# Generation and MHC class II loading of an endogenous influenza epitope revealed by a T cell receptor-like antibody

**DOI:** 10.64898/2026.06.25.734564

**Authors:** Linlin Yang, Michael J. Hogan, Stephen D. Carro, Emma J. Hedgepeth, Jianqiu Du, Mary E. O’Mara, Kathleen S. Krauss, Robert Novak, Christabel Ameyaw Baah, Nhu Le, Chaitali Bhadiadra, Jesper Pallesen, Laurence C. Eisenlohr

## Abstract

CD4^+^ T cells coordinate immune responses to infections and cancers and initiate many autoimmune diseases, yet the intracellular pathways that generate the peptide antigens they recognize remain incompletely understood. Progress has been limited in part by the scarcity of reagents that directly detect defined peptide:MHC class II (pMHC-II) complexes. Here, we established an mRNA vaccine-based pipeline to generate and characterize T cell receptor (TCR)-like antibodies. One result was 1D6, a monoclonal antibody specific for NA25:A^b^, an influenza A/PR8 neuraminidase-derived epitope presented by MHC-II. 1D6 bound NA25:A^b^ with high affinity and specificity. 1D6 recognizes NA25:A^b^ through binding determinants partially distinct from those used by the cognate TCR. Using this reagent, we confirmed the proteasome dependence of NA25 presentation and showed that NA25:A^b^ accumulates in intracellular MHC-II loading compartments without a major requirement for canonical macroautophagy, pointing to an unconventional mode of antigen presentation. These findings establish TCR-like antibodies as powerful tools for dissecting noncanonical MHC-II antigen processing pathways.

**One Sentence Summary:** An mRNA immunization strategy generated a highly specific antibody to study how a noncanonical natural epitope from influenza virus is produced and loaded onto MHC-II.

## INTRODUCTION

By convention, CD4^+^ T cells (T_CD4+_) are activated by MHC class II (MHC-II)-bound peptides (“epitopes”) derived from extracellular antigens (*1*). However, we and many others have demonstrated that MHC-II-bound peptides can originate from intracellular (“endogenous”) proteins encoded either by the cell itself (*2–5*), the genomes of infecting viruses (*6–8*), or even synthetic mRNA delivered by lipid nanoparticles (*9*). Indeed, we have reported that the majority of the T_CD4+_ response to influenza A virus (IAV) is driven by endogenously derived peptides (*10*). Many intracellular pathways have been invoked to explain the conversion (“processing”) of endogenous proteins to MHC-II bound peptide, including macroautophagy (*11*), chaperone-mediated autophagy (*12*), and pathways more associated with MHC class I-(MHC-I)-restricted antigen processing, including dependency on the proteasome and transporter associated with antigen processing (TAP) (*8*, *10*, *13*, *14*). Likewise, many intracellular locations for peptide loading onto MHC-II have been suggested, including the late endosome/lysosome (*15–17*), early endosome (*13*, *18–20*), and endoplasmic reticulum (*21–23*).

To date, most studies of antigen processing and peptide loading, for both MHC-I and MHC-II, have been limited by the standard methods used to detect peptide:MHC (pMHC) complexes. Typically, the readout is T cell activation, which is dependent upon the presence of cognate p:MHC complexes at the surface of the antigen-presenting cell (APC). Information on peptide generation and loading is deduced from the use of gene knockout/knockdown or inhibitors that target specific cellular components. For example, a requirement for expression of the DM chaperone in MHC-II peptide presentation implies loading of peptide in the late endosome, the primary location of this protein (*24*). Another approach for deducing site(s) of peptide loading has been analysis of subcellular fractions for the presence of “SDS-stable” (peptide loaded) MHC-II (*15–17*, *25*), although this approach does not provide information about the loading of specific peptide sequences, which, as mentioned above, can vary depending on the epitope. Loading compartments can also be analyzed by probing subcellular fractions with peptide-specific T cells (*26*) (a technique that proved intractable for us), or, potentially, by mass spectrometric analysis (*27*).

TCR-like antibodies, i.e. those recognizing a specific peptide:MHC-II (pMHC-II) complex, overcome many of these limitations, although the small number of such antibodies reflects the current technical challenges associated with making them (*28*, *29*). Additionally, to our knowledge, no MHC-II-specific TCR-like antibody reported to date detects a pathogen-derived epitope that is generated exclusively, or even mainly through endogenous antigen-processing pathways (*30–33*). To enable studies on the poorly understood cell biology of endogenous MHC-II presentation, we set out to develop a TCR-like monoclonal antibody (mAb) specifically against an endogenously presented viral pMHC-II complex.

Here, we report on the production of a novel TCR-like antibody, 1D6, which is specific for an MHC-II-bound, IAV-derived peptide (NA_97-111_), that is generated only from endogenously expressed source protein. After confirming the specificity of 1D6, we used this antibody to further delineate the antigen processing and presentation pathway(s) used for NA_97-111_, including, most notably, direct identification of the intracellular site of peptide loading.

## RESULTS

### TCR-like mAb clone 1D6 recognizes NA25:A^b^ complexes on flu-infected cells

To enable studies that elucidate endogenous MHC-II presentation pathways, we developed a TCR-like mAb recognizing an endogenously presented neuraminidase (NA)-derived peptide, NA_97-111_, from IAV strain A/Puerto Rico/8/1934 (PR8) that can be presented on the mouse C57Bl/6 MHC-II molecule, H2-A^b^ (A^b^). We refer to this peptide as NA25 based on its numbering in the PR8 NA peptide array available from BEI Resources. We previously showed that NA25 presentation was strongly, if not exclusively, dependent upon translation of NA protein within APCs and that it could be abrogated by treating APCs with an inhibitor of the proteasome (*10*). Thus, NA25 presentation uses non-classical antigen presentation pathways, while its exact presentation pathway remains poorly elucidated.

A TCR-like mAb against NA25:A^b^ was generated as depicted in Fig. 1A. As an immunogen, we designed an mRNA-encoded single chain MHC-II construct where the NA25 peptide was genetically fused to the β chain of A^b^ (A^b^β), and the α chain of A^b^ (A^b^α) was expressed on a separate, co-delivered mRNA to allow formation of full NA25:A^b^ complexes. BALB/c (MHC-mismatched) mice were immunized with this construct in the form of a nucleoside-modified mRNA-liposome vaccine (*34*). We screened mouse sera for IAV reactivity by first adsorbing out non-NA25-dependent A^b^ reactivity, which was dominant given the MHC mismatch, and then used the adsorbed sera to stain IAV-infected vs. uninfected fibroblasts. All three immunized mice showed specific reactivity of adsorbed serum to flu-infected MHC-II^+^ cells (Fig. 1B). Splenocytes were stained with NA25:A^b^ tetramers conjugated to two different fluorophores, and double tetramer^+^ cells were singly sorted (Fig. 1C, fig. S1) onto monolayers of NB-21 feeder cells, which cause B cells to proliferate and secrete IgG (*35*, *36*). While numerous supernatants showed IgG binding to NA25:A^b^ protein by ELISA, only 4 out of 213 clones showed notable binding to flu-infected MHC-II^+^ cells (Fig. 1D). Of these, we observed consistent specific binding with supernatant from one clone, 1D6 (fig. S2A), which was selected for B cell receptor cloning and recombinant expression. We confirmed that recombinant 1D6 strongly stained IAV-infected but not uninfected mouse cells and that this was strictly dependent on A^b^ expression (Fig. 1E). In line with this finding, we found that the MHC-II-blocking antibody M5/114 abrogated 1D6 recognition of IAV PR8-infected A^b+^ cells (fig. S2B). 1D6 also strongly stained A^b+^ cells that were pre-incubated with synthetic, soluble NA25 peptide but not an irrelevant IAV peptide (Fig. 1F). 1D6 exhibits some level of strain specificity, as it recognized MHC-II^+^ cells infected with IAV strain PR8 but not X31 (Fig. 1G, fig. S2A), which encodes a different subtype of NA protein (*37*). We further demonstrated that 1D6 could inhibit activation of an NA25:A^b^-specific T_CD4+_ hybridoma (*10*) in a co-culture with IAV-infected cells (Fig. 1H). As expected, given the dependence of NA25 epitope generation on endogenously derived NA protein, 1D6 staining of flu-treated MHC-II^+^ fibroblasts was restricted to NA^+^ infected cells, with no detectable staining in NA^-^ bystander cells (Fig. 1I). Notably, 1D6 was able to detect NA25:A^b^ presentation by IAV-infected cells with a similar level of sensitivity as the hybridoma (Fig. 1J).

**Fig. 1.**
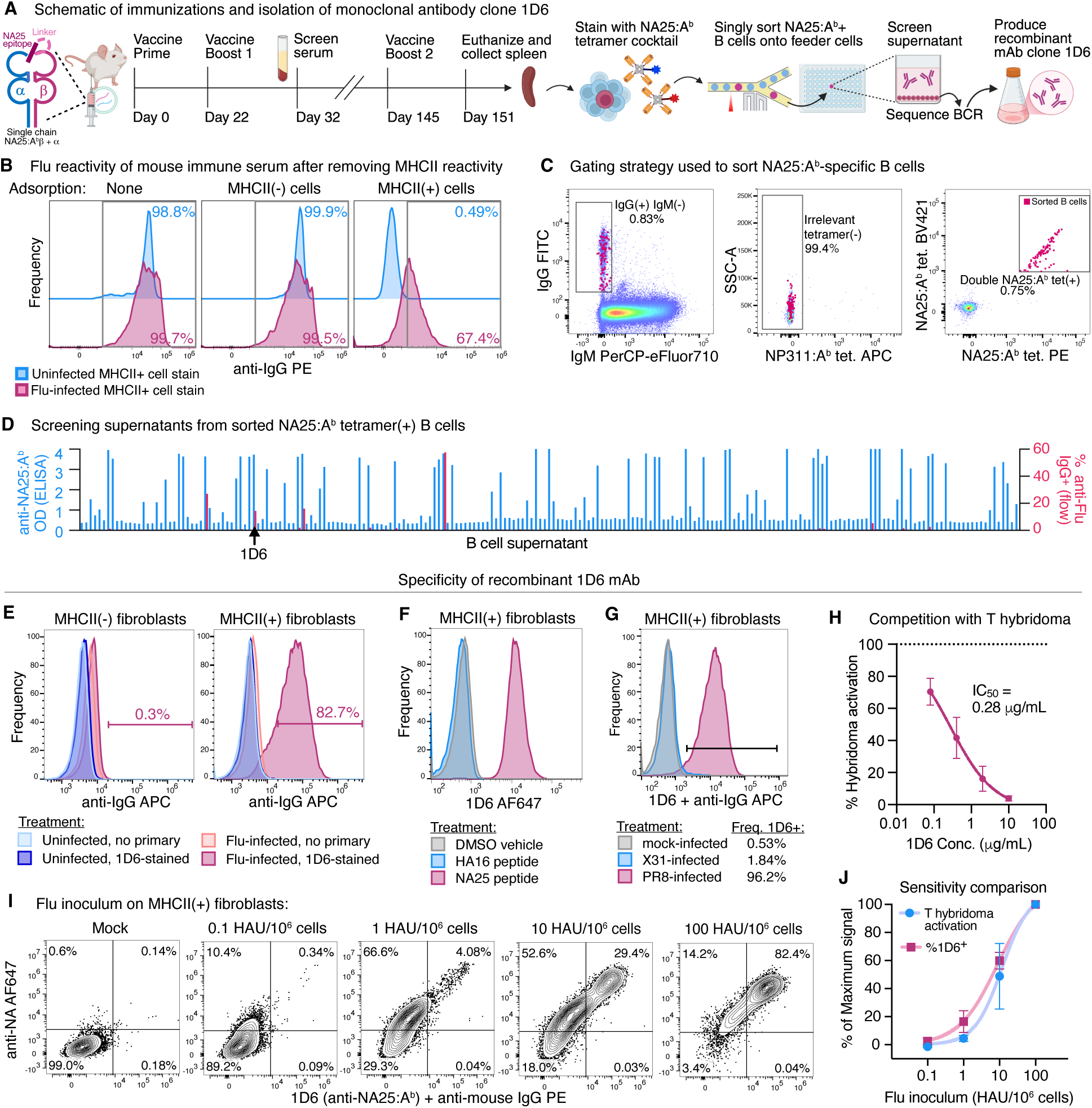
Generation and characterization of a TCR-like mAb, 1D6, recognizing IAV PR8 NA25:A^b^ complexes. (**A**) Timeline and schematic of immunizations and B cell receptor isolation. (**B**) IAV-dependent reactivity of mouse immune serum was measured by first adsorbing out general A^b^ reactivity from serum, then staining IAV PR8-infected vs. uninfected A^b+^ cells (B6-CIITA-E^d^ fibroblasts) with the adsorbed serum. (**C**) Simplified gating strategy used to singly sort NA25:A^b^-specific B cells onto NB-21 monolayers. (**D**) Supernatants from 213 NB-21/B cell co-cultures from selected mouse were screened by ELISA (left y-axis, blue bars) for IgG binding of monomeric NA25:A^b^ protein and by flow cytometry (right y-axis, red bars) for IgG binding of IAV PR8-infected B6-CIITA-E^d^ fibroblasts. (**E**) Recombinant, purified 1D6 mAb was validated for specific staining of MHCII(+) B6-CIITA-E^d^ fibroblasts (right) and not MHCII(-) parental B6 fibroblasts (left) infected with IAV PR8 at 50 HAU per million cells. (**F**) 1D6 binding to cells pulsed with synthetic NA25 peptide but not irrelevant HA16 peptide. (**G**) 1D6 binding to B6-CIITA-E^d^ cells infected overnight at MOI of 1 with IAV PR8 (H1N1) but not X31 (H3N2). (**H**) Competition between 1D6 binding and NA25-specific T_CD4+_ hybridoma activation by PR8-infected B6-CIITA-E^d^ cells. (**I**) Comparison of 1D6 and anti-NA mAb binding to B6-CIITA-E^d^ cells infected overnight with PR8. (**J**) Comparison of sensitivity of 1D6 staining vs. NA25-specific T_CD4+_ hybridoma activation by B6-CIITA-E^d^ cells infected overnight with the indicated amounts of PR8.

### 1D6 recognizes NA25:A^b^ complex differently from a native TCR

MHC class II molecules accommodate peptide ligands within an open-ended groove-like binding site (*38*), such that – in general – amino acid residues at positions P2, P3, P5, P7, and P8, as well as those flanking the core P1-P9 binding register, remain solvent exposed and therefore accessible for TCR contact (*39–41*). To investigate if our TCR-like antibody engages the pMHC-II complex similarly to an NA25:A^b^-specific TCR, we used a panel of truncated peptides for epitope mapping to define the core binding determinants of 1D6. To compare antibody binding to TCR recognition, 1D6 binding was measured alongside the activation of our NA25-specific T_CD4+_ hybridoma cell line. The binding of 1D6 was highly dependent on the N- and C-terminal core epitope flanking residues, but the C-terminal histidine was more critical for binding than the N-terminal glycine (Fig. 2A). In contrast, TCR recognition of NA25 was much more tolerant of single residue deletions at the peptide termini, with no impact on hybridoma activation (Fig. 2A). Furthermore, TCR recognition was not fully abrogated until removal of both terminal and two internal residues. Together, these data suggest somewhat distinct modes of NA25:A^b^ complex recognition between the 1D6 antibody and this native TCR (Fig. 2B).

**Fig. 2.**
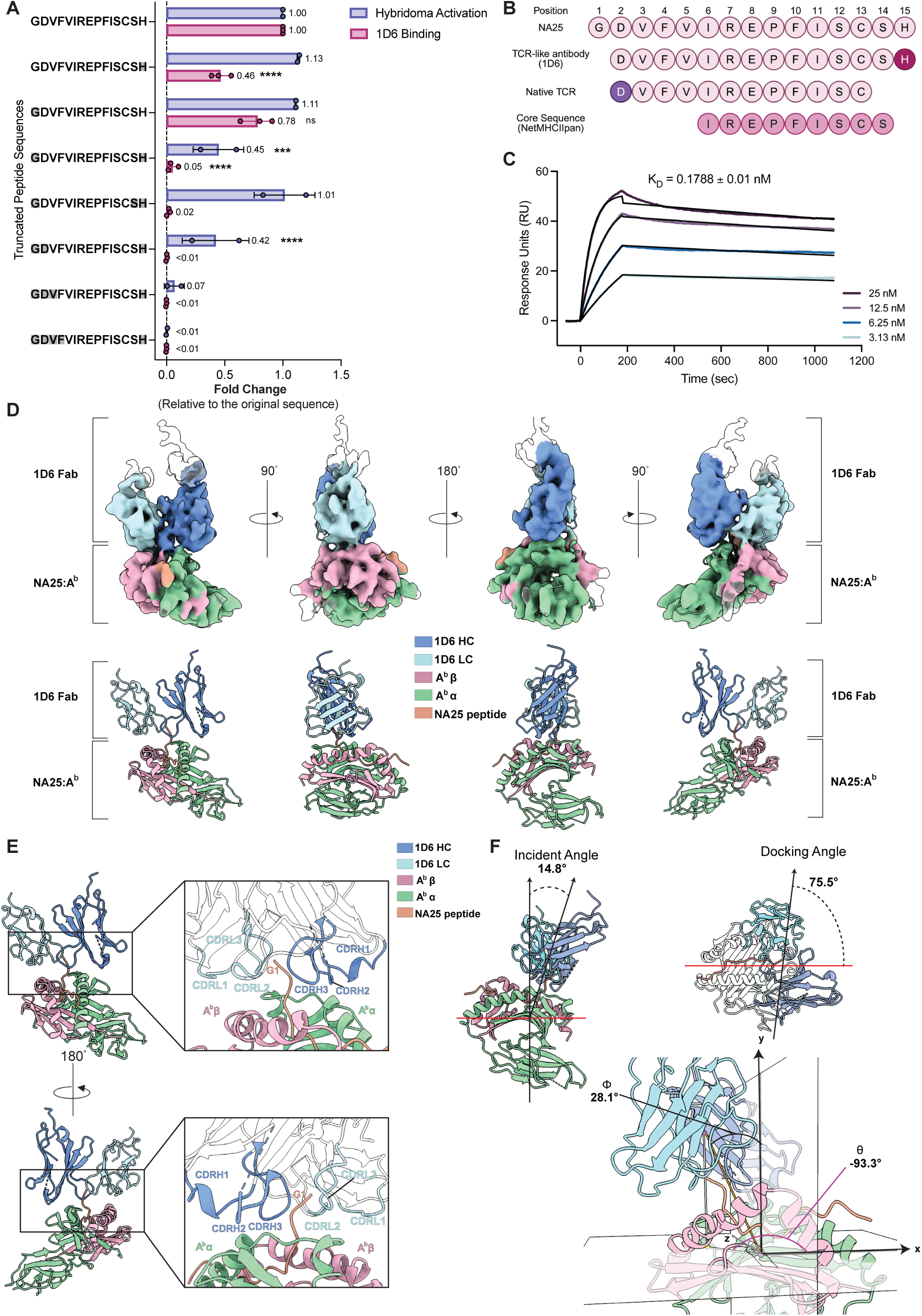
Binding characterization of 1D6 and structural basis of 1D6 Fab recognition of NA25:A^b^. (**A**) Epitope mapping of 1D6 compared with cognate T_CD4+_ hybridoma activation. A panel of NA25 peptides with progressive truncations from the N- or C-terminus (truncated residues indicated in gray) was assessed for 1D6 binding (flow cytometry) or activation of NA25-specific T_CD4+_ hybridoma. Signals were normalized to “full-length” NA25 (NA_97–111_), which provides maximal signal in both cases. Data represent the combined results of two (hybridoma activation) or three (1D6 binding) independent biological experiments. (**B**) Alignment of residues required for 1D6 binding and cognate TCR activation mapped onto the NA25 sequence. The predicted MHC class II core binding register [NetMHCIIpan-4.0 (*120*)) is highlighted. This 9-mer binding core is predicted to directly interact with the MHC binding groove. (**C**) Representative surface plasmon resonance (SPR) sensorgrams showing binding of 1D6 to immobilized biotinylated NA25:A^b^ monomer on a streptavidin chip across a range of analyte concentrations (indicated). Black lines represent global kinetic fits. Data are presented as mean ± s.d. from n = 2 technical replicates. (**D**) Cryo-EM density map and refined model of the 1D6/NA25:A^b^ complex. Each component is colored as indicated. (**E**) Interface analysis of 1D6 CDR loops with NA25:A^b^ represented in two views. (**F**) Binding geometry analysis of 1D6 with NA25:A^b^. The upper panel shows conical angle calculation. The bottom panel shows TCR CoM spherical parameters. (**A**) Two-way ANOVA with Šídák’s multiple comparisons test compared to full length signals.

### Biophysical and structural characterization of the 1D6 Fab/NA25:A^b^ complex

TCRs generally have low affinity for ligands (dissociation constant [K_D_] ∼1-100 µM), a feature thought to facilitate rapid sampling of pMHC complexes during antigen surveillance (*42–45*). Since our epitope-mapping data suggested that 1D6 recognizes NA25:A^b^ differently from the cognate TCR, we hypothesized that this distinct pattern of peptide engagement might also be reflected in an atypical binding affinity and structural mode of pMHC-II recognition. To first quantify the affinity of 1D6 to NA25:A^b^ monomer, we performed surface plasmon resonance (SPR), using the irrelevant NP47:A^b^ monomer as a negative control. 1D6 bound NA25:A^b^ monomer with high affinity (K_D_=1.788×10^-10^ M) (Fig. 2C), whereas no detectable binding to NP47:A^b^ was observed (fig. S2C). Thus, 1D6 binds to its cognate pMHC-II complex with high specificity and affinity.

We next sought to determine the structural basis for binding of NA25:A^b^ by 1D6 (Fig. 2, A and B). To facilitate our structural characterization, we produced and purified peptide-linked NA25:A^b^ monomer, where the NA25 peptide was covalently bound to the A^b^β chain via a glycine-serine linker. To promote heterodimer formation and stabilize pairing, each MHC-II chain was fused to a heterodimeric coiled coil motif derived from the leucine zipper proteins Fos and Jun, as previously described (*46*) (fig. S2, D-F).

To understand the engagement of 1D6 to NA25:A^b^, we formed the 1D6 Fab/NA25:A^b^ complex and obtained a cryo-EM density map at 5.1 Å resolution. We docked the models of 1D6 variable regions (VH/VL) (generated by SWISS-MODEL) (*47*) and NA25:A^b^ (PDB 6MNG) (Fig. 2D, fig.S2G and Table S1). By refining the Cα backbone of CDR loops of 1D6 Fab, we observed that the CDRL3 of the light chain and the CDRH3 of the heavy chain interact with the N-terminal region of the NA25 peptide, completely surrounding the N-terminal region in a clamp-like fashion (Fig. 2E). This interaction pulls the N-terminal region away from the peptide-binding groove. Furthermore, the CDRL1 and CDRL2 interact with the A^b^β chain whereas CDRH1 and CDRH2 interact with the A^b^α chain (Fig. 2E).

Next, we analyzed the binding geometry of 1D6 Fab/NA25:A^b^ complex using two complementary approaches (*43*, *48*). We first used conical-angle calculations to determine the orientation of 1D6 relative to the NA25:A^b^ peptide-binding groove (*43*). This analysis showed an incident angle of approximately 15°, and a docking angle of approximately 76° (Fig. 2F, Top). We also calculated the spherical parameters using TCR CoM, which describes the center-of-mass (CoM) position of the TCR, or in our case the TCR-like antibody, relative to the MHC CoM (*48*). This analysis yielded a θ angle of ∼-93° and a ϕ angle of ∼28° (Fig. 2F, Bottom). When comparing with currently available TCR/pMHC-II structural data, only two TCR/pMHC-II complexes, PDB 2WBJ and 1YMM, showed docking angles close to that observed for 1D6. The incident and docking angles for PDB 2WBJ and 1YMM are 20°, 80° and 18°, 84°, respectively. Notably, among the analyzed TCR/pMHC-II complexes, no TCR exhibited a docking angle between 60° to 80° (*49*). Thus, 1D6 engages NA25:A^b^ in a similar but rather unique manner in terms of peptide engagement as well as docking angle compared to other previously published, naturally occurring TCRs.

### Endogenous MHC-II-restricted antigen processing and presentation revealed by 1D6

A major application of TCR-like antibodies is to probe the intracellular processing pathways that contribute to antigen presentation. Using T_CD4+_ hybridoma-based assays, we have previously described several requirements for the presentation of NA25 from PR8 (*10*). Due to the difference between TCR and 1D6 engagement of NA25:A^b^ complexes, we asked whether our previous findings using epitope-specific T_CD4+_ hybridoma assays could be recapitulated with 1D6. First, we have demonstrated that NA25 is an endogenous epitope whose processing and presentation requires synthesis of the parent antigen within the APC (*10*). To reaffirm the requirement for endogenous parent protein, we treated APCs with increasing doses of infectious (“live”) PR8 or chemically inactivated PR8. As expected, we observed hybridoma activation from live PR8 but not from inactivated PR8 (fig. S3A) despite abundant NA protein in similar conditions (*50*). Consistent with this, 1D6 staining was absent on cells expose to inactivated PR8 even at a dose equivalent to a multiplicity of infection (MOI) of 200, while infectious PR8 generated a robust NA25:A^b^ signal at an MOI of 5 (Fig. 3A). We thus confirmed that NA25 is a strongly *endogenous* MHC class II-restricted influenza epitope.

**Fig. 3.**
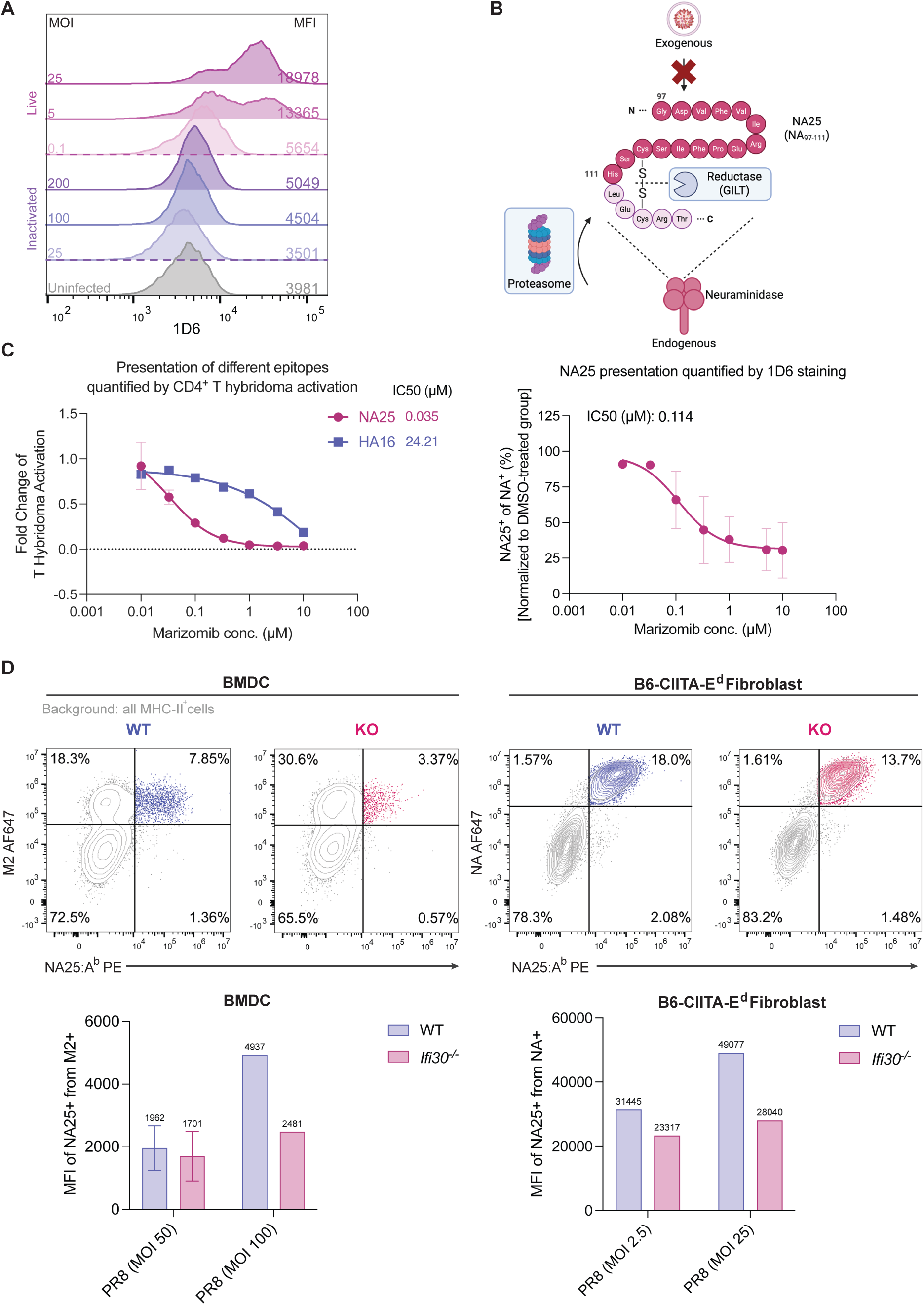
Endogenous MHC-II-restricted antigen processing and presentation revealed by 1D6. (**A**) B6-CIITA-E^d^ fibroblasts were infected with live PR8 (MOI 0.1, 5, and 25) or β-Propiolactone (BPL)-inactivated PR8 (at doses equivalent to MOI 25, 100, and 200) for 16-18 hours. Intracellular and surface NA25:A^b^ complexes were detected using 1D6 antibody. (**B**) Schematic of intracellular components that have been previously reported to be involved in NA25 epitope endogenous processing and presentation (*46*). (**C**) (Left) NA25- and HA16-specific T_CD4+_ hybridoma were used to assess the effect of marizomib on epitope presentation following PR8 infection. Results are representative of three independent experiments. (Right) 1D6 staining quantifying NA25:A^b^ presentation across increasing concentrations of marizomib following PR8 infection. Data from four independent experiments were pooled. IC_50_ values were determined using a sigmoidal 4PL model. (**D**) Cells were infected with PR8 at MOI 100 (BMDCs) or 2.5 (fibroblasts). M2: IAV Matrix-2 protein, an ion channel protein. Representative flow plots are shown from three independent experiments (BMDCs). MFI values for NA25^+^ cells within the M2^+^ or NA^+^ population were pooled from independent experiments across cell types and PR8 MOIs.

We previously demonstrated that NA25:A^b^ presentation requires intracellular components that are more associated with MHC-I presentation than classical MHC-II antigen presentation, including the proteasome (*10*) (Fig. 3B). Therefore, we investigated the impact of proteasome inhibition on 1D6 staining by treating APCs with marizomib, an irreversible pan-proteasome inhibitor that targets all three proteolytic activities of the 20S proteasome (*51–53*). In line with our previous finding (*10*), we observed that the presentation of NA25, measured via hybridoma activation or 1D6 binding, is highly sensitive to marizomib treatment, with decreasing signal in a dose-dependent manner (Fig. 3C). Notably, presentation to a T_CD4+_ hybridoma clone specific for a PR8 hemagglutinin-derived epitope, HA16, which can be processed via both endogenous and exogenous pathways (*10*), exhibited reduced sensitivity to marizomib treatment, with a ∼700-fold greater IC_50_ (Fig. 3C). Together, these data demonstrate that 1D6 binding at the cell surface is dependent on the proteasome and affirm the critical role of the proteasome in generating NA25. Furthermore, 1D6 staining provides the advantage of being able to quantify the percentage of NA25:A^b+^ cells out of the infected (NA^+^) population (Fig. 3C), providing a more precise assessment than which can be inferred from the hybridoma assay.

Previously, we reported that γ-interferon–induced lysosomal thiol reductase (GILT, encoded by *Ifi30* in mice), involved in the reduction of many disulfide-bonded antigens (*54*), is critical for the processing and presentation of endogenous NA25 in bone marrow-derived dendritic cells (BMDCs) (*10*). In line with this, homology-based structural analysis (see Methods) indicates that generation of the NA25 epitope from native neuraminidase may be constrained by a putative Cys109-Cys114 disulfide bond. To correlate previous findings with the T_CD4+_ hybridoma of NA25:Ab specificity (*10*), we investigated 1D6 staining in *Ifi30^-/-^* BMDCs (fig. S3B) and fibroblasts (fig. S3C). In *Ifi30^-/-^* BMDCs, 1D6 staining was reduced both in percentage as well as mean fluorescence intensity (MFI) (Fig. 3D), consistent with the previously established GILT dependency of NA25 presentation. Likewise, optimal 1D6 staining in fibroblasts also required GILT expression (Fig. 3D and fig. S3D), in agreement with NA25 presentation measured via hybridoma activation assay (fig. S3E). Collectively, these findings demonstrate the utility of 1D6 in the investigation of processing pathways that drive presentation of NA25 and further highlight its strong concordance with epitope-specific T_CD4+_ hybridoma assays.

### NA25:A^b^ complexes are localized within LAMP-1+ compartments

A fundamental question of endogenous MHC-II antigen processing and presentation is where pMHC-II complexes are generated intracellularly. While classical pathways place MHC-II peptide loading within the late endosome, the juxtaposed proteasome (cytosol) and GILT (late endosome/lysosome (*55*)) dependence of NA25 leaves the localization of peptide loading hard to predict. To date, there has been limited usage of TCR-like antibodies to address this question. Therefore, we evaluated the capacity for 1D6 to be used for immunofluorescence (IF) staining to localize NA25:A^b^ complexes within the cell.

Adapting methods from previous publications using pMHC-specific antibodies for IF microscopy (*56–58*), we infected B6-CIITA fibroblasts with PR8 and performed IF staining using 1D6 16 hours post-infection. We observed significant 1D6 intracellular staining in PR8-infected but not X31-infected cells (fig. S4A), reaffirming the specificity of 1D6 staining. We next investigated the co-localization of 1D6 stain with markers of subcellular compartments of interest within PR8-infected cells. The Y-Ae antibody, which detects the Eα_52-68_:A^b^ complex derived from the α chain of H2-E^d^ (*31*), has been previously shown to stain LAMP-1+ compartments (i.e., late endosome/lysosome) (*59*). We therefore asked whether 1D6 staining similarly localizes in LAMP-1+ compartments, using the cis-Golgi marker GM130 for comparison. We observed significantly greater correlation and co-localization of 1D6 with LAMP-1 signal compared to GM130 (Fig. 4, A-D). These data suggest that NA25:A^b^ complexes localize to classical LAMP-1+ compartments, despite its unconventional, proteasome-dependent antigen processing pathway.

**Fig. 4.**
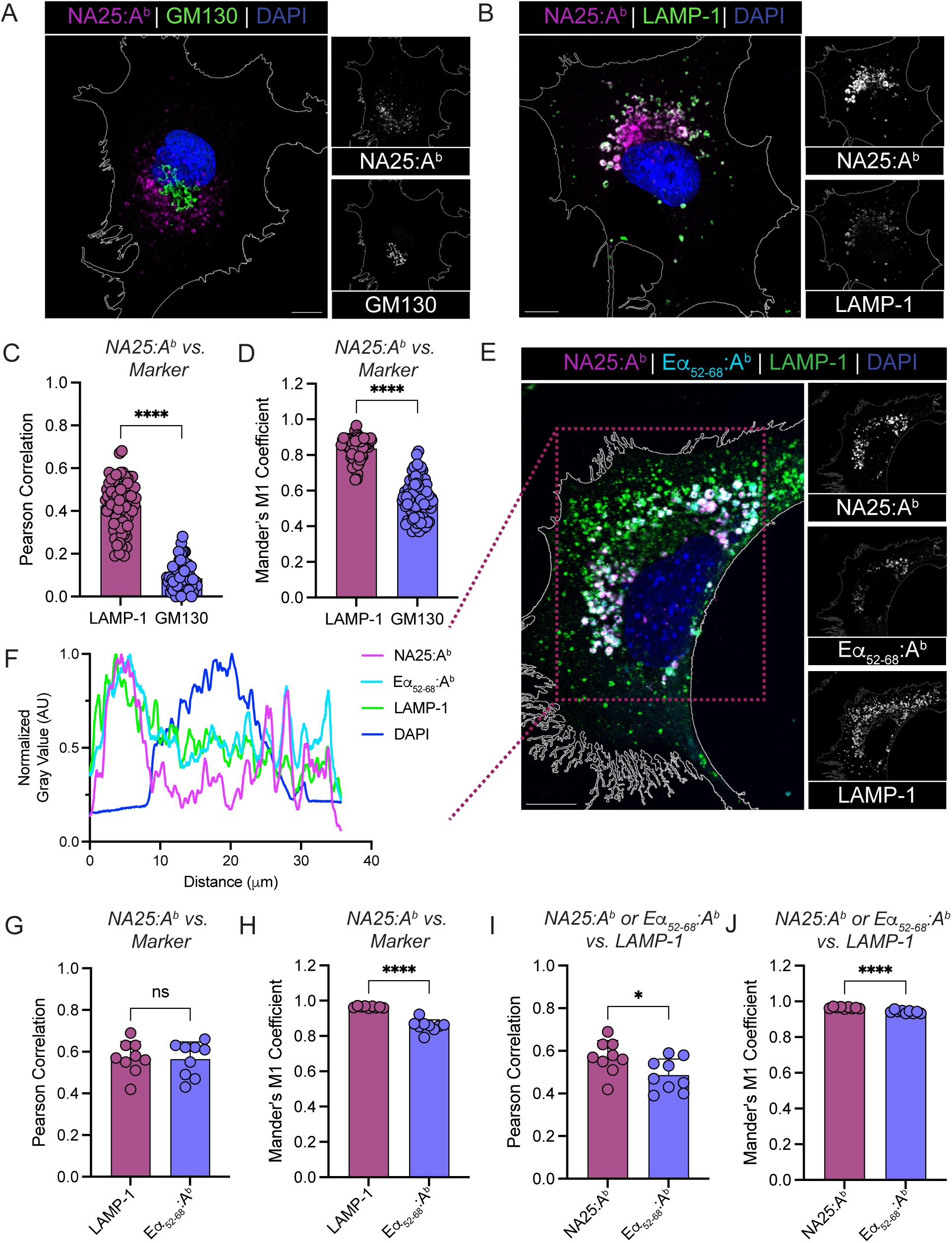
NA25:A^b^ complexes localize to LAMP-1+ compartments. (**A**) Representative composite and single channel immunofluorescent images of NA25:A^b^ complexes (stained with 1D6, magenta) and GM130 (green) in B6-CIITA cells, with DAPI used as a counterstain (blue). Scale bar indicates 10 µm. (**B**) Representative composite and single channel immunofluorescent images of NA25:A^b^ complexes (1D6, magenta) and LAMP-1 (green) in B6-CIITA cells, with DAPI used as a counterstain (blue). Scale bar indicates 10 µm. (**C**) Pearson correlation between NA25:A^b^ and LAMP-1 or GM130 signal in B6-CIITA cells. n = 79 cells for LAMP-1 and n = 75 cells for GM130. (**D**) Mander’s M1 coefficient between NA25:A^b^ and LAMP-1 or GM130 signal in B6-CIITA cells. n = 79 cells for LAMP-1 and n = 75 cells for GM130. (**E**) Representative composite and single channel immunofluorescent images of NA25:A^b^ complexes (1D6, magenta), Eα_52-68_:A^b^ complexes (Y-Ae, cyan), and LAMP-1 (green) in B6-CIITA-E^d^ cells, with DAPI used as a counterstain (blue). Scale bar indicates 10 µm. (**F**) Intensity (grey value) profile of each individual channel along the rectangular boundary overlaid in (E). Mean grey values at each distance were normalized to the maximal signal in each corresponding channel. (**G**) Pearson correlation between NA25:A^b^ and LAMP-1 or Eα_52-68_:A^b^ signal in B6-CIITA-E^d^ cells. n = 10 cells. (**H**) Mander’s M1 coefficient between NA25:A^b^ and LAMP-1 or Eα_52-68_:A^b^ signal in B6-CIITA-E^d^ cells. n = 10 cells. (**I**) Pearson correlation between NA25:A^b^ or Eα_52-68_:A^b^ and LAMP-1 signal in B6-CIITA-E^d^ cells. n = 10 cells. (**J**) Mander’s M1 coefficient between NA25:A^b^ or Eα_52-68_:A^b^ and LAMP-1 signal in B6-CIITA-E^d^ cells. n = 10 cells. (**C**-**D**, **G**-**J**) Unpaired t-test.

To directly compare 1D6 signal with Y-Ae, we infected B6-CIITA-E^d^ cells with PR8 and co-stained for 1D6, Y-Ae, and LAMP-1. Notably, we observed similar spatial distributions of 1D6, Y-Ae, and LAMP-1 signals within the cell (Fig. 4, E and F and fig. S4B). Furthermore, 1D6 signal was equally correlated with LAMP-1 or Y-Ae (Fig. 4G) by Pearson coefficient but had significantly greater overlap with LAMP-1 than Y-Ae as measured by Mander’s M1 coefficient (Fig. 4H), suggesting that a greater proportion of 1D6 signal co-localizes with LAMP-1 than with Y-Ae. In line with this, 1D6 signal was more strongly correlated and co-localized with LAMP-1 whereas Y-Ae signal correlated and co-localized less with LAMP-1 (Fig. 4, I and J). Together, our data demonstrate strong colocalization of NA25:A^b^ complexes with LAMP-1+ compartments, in line with its dependence upon GILT expression (previously reported to localize to the late endosome/lysosome (*55*)), for efficient presentation.

### Macroautophagy is not involved in NA25 presentation

Macroautophagy delivers cytosolic cargo to late endosomes and lysosomes and has been reported many times to be a mechanism for endogenous MHC-II processing and presentation (*60–63*), raising the possibility that this pathway contributes to the presentation of NA25. Macroautophagy is initiated by phagophore formation and elongation, which requires highly coordinated activity of kinases, including PI3K complexes (*64*, *65*). This is followed by autophagosome maturation through a ubiquitin-like conjugation cascade that converts LC3-I to LC3-II via conjugation to the lipid phosphatidylethanolamine (PE) (*66*). We and other groups have shown that IAV infection induces functional macroautophagy (*67–69*), although direct evidence linking this process to MHC-II-restricted antigen presentation remains limited (*69*). To inhibit macroautophagy, we used two complementary pharmacologic inhibitors of PI3K-dependent autophagy initiation with distinct selectivity and toxicity profiles. SAR405 selectively inhibits PI3K class III/Vps34 (*70*, *71*), whereas wortmannin broadly inhibits all PI3K family members (*72*), and is associated with greater cytotoxicity. We first treated B6-CIITA-E^d^ fibroblasts with increasing concentrations of SAR405 beginning 1 hour before PR8 infection. To verify inhibition, we assessed LC3-I to LC3-II conversion by western blot. Following treatment with 0.5 µM SAR405, we observed accumulation of LC3-I relative to the DMSO-treated control, consistent with impaired LC3 processing (Fig. 5A). LC3-II levels did not show obvious reduction, potentially reflecting continued autophagy induction during viral infection (Fig. 5A). At the concentrations tested (0.5 µM and 1 µM), SAR405 had no measurable effect on cell viability, infection efficiency, or 1D6 staining among infected cells (NA^+^) (Fig. 5B). To validate that SAR405 does impair macroautophagy-dependent MHC-II antigen presentation in our system, we established a functional positive-control assay, as previously described (*69*). We fused the classically presented E^d^-restricted influenza hemagglutinin Site-1 (S1) epitope, which has intrinsic resistance to endocytic proteases (*19*, *73*), to the N-terminus of LC3B, a protein known to localize to autophagosomes (*61*, *74*), and measured S1 presentation using S1-specific T_CD4+_ hybridomas with or without SAR405 treatment (fig. S5A). In parallel, we generated an LC3B G120A mutant, which prevents LC3 lipidation and membrane conjugation (*66*, *69*). Twenty-four hours after plasmid transfection, cells were acid-stripped with glycine to remove preexisting surface pMHC-II complexes, cultured overnight in the presence of SAR405, and fixed prior to co-culture with T_CD4+_ hybridomas. In contrast to what we found with 1D6 staining, SAR405 treatment significantly reduced S1 presentation in cells expressing wild-type S1-LC3B but not in cells expressing the lipidation-defective S1-LC3B (G120A) (fig. S5B). Together, these results confirm the biological activity of SAR405 under our experimental conditions and indicate that 1D6 staining is insensitive to macroautophagy inhibition via SAR405 treatment.

**Fig. 5.**
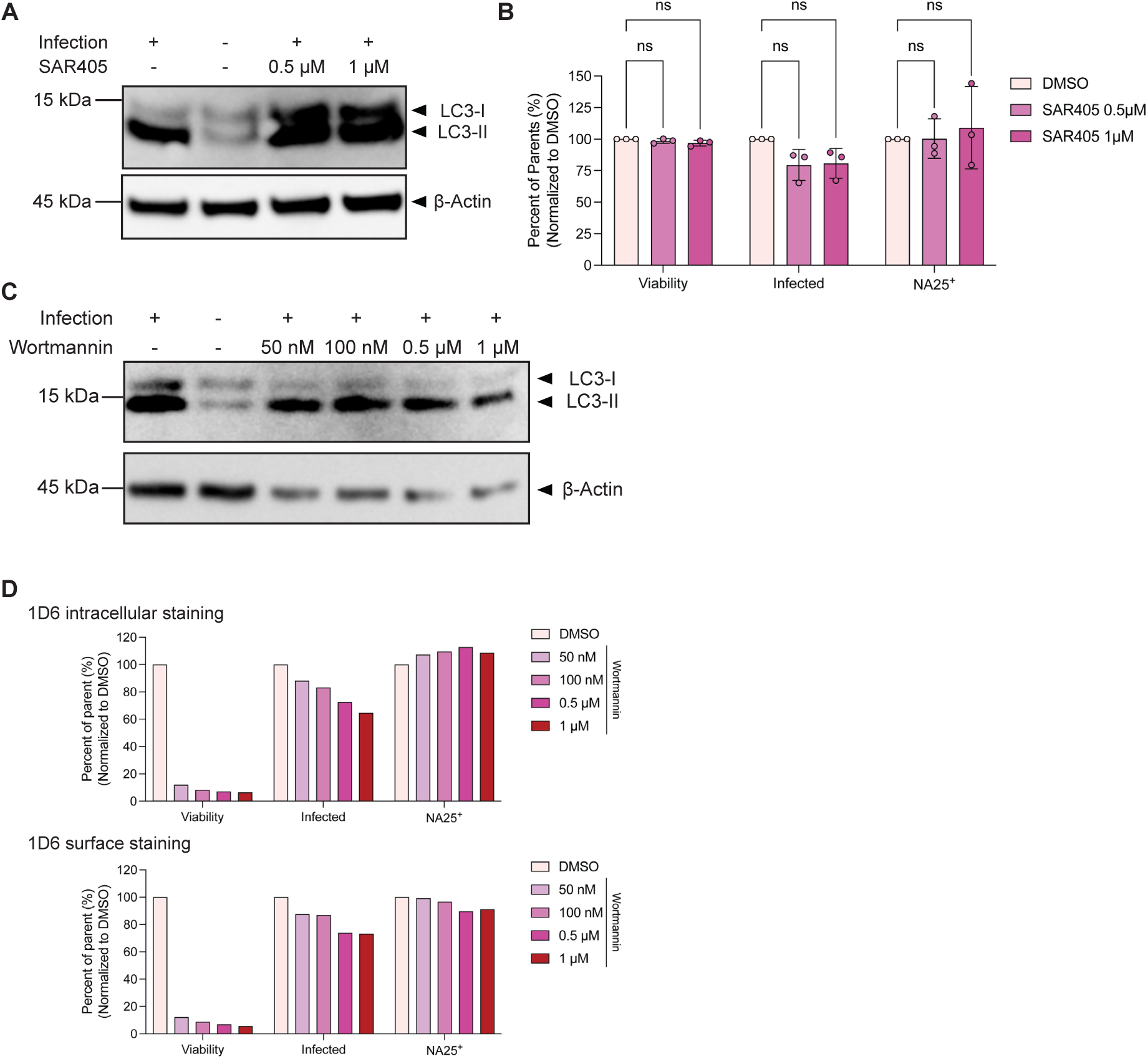
Macroautophagy is not involved in the NA25 presentation. (**A**) and (**C**) Immunoblot analysis of LC3 conversion in lysates from B6-CIITA-E^d^ fibroblasts infected with PR8 (MOI 10) and treated with SAR405 (**A**) or wortmannin (**C**) at the indicated concentrations. Uninfected cells without inhibitor treatment were included to establish the baseline expression of LC3-I and LC3-II. β-actin served as a loading control. (**B**) Cell viability, infectivity (percentage of NA^+^ cells within the live population), and NA25:A^b+^ among NA^+^ cells under the indicated treatments. Data were pooled from three independent flow experiments and normalized to the DMSO-treated condition across experiments. Non-significant differences were determined by two-way ANOVA with multiple comparisons versus DMSO. (**D**) 1D6 staining of PR8-infected cells 16 to 18 hours after infection. Top, intracellular staining; cells were fixed and permeabilized before staining. Bottom, surface staining only; cells were fixed without permeabilization before staining.

As an orthogonal approach, we next treated cells with wortmannin and similarly assessed impact on NA25 presentation during PR8 infection via 1D6 staining. Wortmannin treatment altered LC3-I/LC3-II patterns during PR8 infection; however, β-actin signal was reduced in wortmannin-treated samples (Fig. 5C), consistent with the increased cytotoxicity observed under these conditions (Fig. 5D). Despite its cytotoxicity, wortmannin did not reduce 1D6 staining among live, infected cells, as measured by the frequency of NA25:A^b^-positive cells within the NA^+^ population (Fig. 5D). Together, these data suggest that canonical macroautophagy does not play a major role in NA25 processing and presentation under these fixed conditions, while further highlighting the utility of this TCR-like antibody as a sensitive, scalable, and experimentally tractable tool that enables straightforward quantification of antigen processing and presentation pathways beyond the resolution of traditional T_CD4+_ hybridoma assays.

## DISCUSSION

Extensive efforts have been directed toward generating antibodies that recognize pMHC-II complexes (*32*, *33*, *75–79*), yet such reagents remain rare, particularly for naturally processed pathogen-derived epitopes. This limitation reflects both the structural properties of MHC-II (*80–82*) and the biological complexity of MHC-II antigen processing, in which peptide length, binding register, and proteolytic trimming may differ depending on the cellular context (*83–85*). Here, we used a single-chain pMHC-II mRNA immunization strategy to generate 1D6, a TCR-like antibody that specifically recognizes the influenza-derived NA25:A^b^ complex with high affinity but in a mode distinct from endogenous TCR. After extensive validation, we used 1D6 to directly detect NA25:A^b^ presentation by infected cells and to reassess key features of NA25 processing that had previously been inferred from T_CD4+_ hybridoma activation assays. Consistent with prior findings, 1D6 confirmed that NA25 presentation is predominantly derived from an endogenous source and depends on proteasomal activity and GILT expression (*10*), thereby affirming the unique and non-classical processing pathway that leads to NA25 presentation by MHC-II. Staining with 1D6 further revealed that intracellular NA25:A^b^ localizes to LAMP-1+ compartments, with pharmacologic inhibition of macroautophagy not diminishing NA25:A^b^ presentation. These findings establish 1D6 as a reagent for direct visualization and quantification of a naturally processed, endogenous MHC-II–restricted viral epitope.

A central advantage of 1D6 is its ability to detect naturally processed NA25:A^b^ complexes generated during influenza infection. Although 1D6 also recognizes synthetic peptide-loaded NA25:A^b^, its recognition of infection-derived complexes enables measurement of this endogenous viral epitope in the cellular context in which it is produced. Unlike T_CD4+_ hybridoma activation assays, which provide a functional readout of T cell stimulation, 1D6 staining permits direct quantification of NA25:A^b^ abundance at the single-cell level and within defined infected cell populations. Thus, 1D6 provides a more precise framework for dissecting how antigen expression, infection efficiency, cellular state, and intracellular trafficking shape endogenous MHC-II presentation. These qualities will be especially useful to interrogate NA25 presentation and the role of key APCs during infection *in vivo*.

The high affinity and TCR-blocking activity of 1D6, combined with our structural data, suggest that this antibody engages NA25:A^b^ in a manner that overlaps with, but is distinct from cognate TCR recognition. Our cryo-EM analysis showed that 1D6 docks over the MHC-II groove in a TCR-like orientation, contacting both peptide and MHC-II residues, but its binding was more sensitive to the removal of epitope flanking residues than our hybridoma (Figure 2). 1D6 utilizes its CDRL3 and CDRH3 loops to surround the N-terminus of NA25 peptide in a clamp-like manner, whereas its CDRL1, CDRL2, CDRH1 and CDRH3 interact with A^b^. This interaction patten is consistent with other reported modes of TCR recognition of pMHC-II complex (*41*, *86*, *87*); however, the clamp-like interaction with the NA25 terminus is quite distinct from canonical TCR/pMHC-II interactions (*40*, *41*). Our structural analysis further reveals that 1D6 engages NA25:A^b^ in a way that differs from most naturally occurring TCRs, exhibiting a distinct docking angle and orientation across the pMHC-II interface. These structural insights may inform future strategies for generating antibodies against naturally processed pMHC-II complexes in the context of infection or autoimmune responses.

Our immunofluorescence imaging with 1D6 indicated that NA25:A^b^ complexes primarily associate with LAMP-1+ compartments (Figure 4), which are classically associated with endolysosomal MHC-II loading during exogenous antigen presentation (*88*). The critical dependence of NA25 presentation on H2-DMα (*10*), which is localized to classical MHC-II loading compartments (*89*), strongly aligns with loading of NA25 onto MHC-II within endolysosomal compartments rather than in alternative secretory organelles, such as the endoplasmic reticulum (*21*). However, the critical dependency of NA25 presentation on the proteasome raises key questions regarding how these two traits are interconnected within the cell. One possibility is that proteasomal processing may occur within, or in close association with, vesicular compartments. A limited number of studies has reported active proteasomes within phagosomal or endolysosomal compartments and proposed that these proteasomes can support TAP-independent, proteasome-dependent cross-presentation (*90*, *91*). Alternatively, NA25 may be generated by cytosolic proteasomes before being delivered to LAMP-1+ MHC-II loading compartments. In this model, neuraminidase could access the cytosol through mechanisms such as retro-translocation (*92*) or mislocalization (*93*), followed by proteasomal processing and subsequent transport of NA25-containing peptide intermediates into endolysosomal compartments. Although NA25 presentation is TAP-independent *in vivo* (*10*), alternative peptide transporters may contribute to this pathway (*94*). TAP-like transporter (TAPL), a TAP homolog localized to lysosomal membranes, has been suggested to transport cytoplasmic polypeptides into the lysosomal lumen (*95*). Autophagy-related pathways may provide another route for delivering cytosolic or cytosol-accessible neuraminidase-derived material into MHC-II loading compartments.

Although macroautophagy has been implicated in delivery of intracellular antigens to MHC-II compartments (*61*, *96–100*), our findings do not support a major requirement for canonical macroautophagy in NA25 presentation. The macroautophagy inhibitors SAR405 and wortmannin modulated LC3 conversion but did not reduce NA25:A^b^ abundance among infected cells, suggesting that NA25 delivery to MHC-II loading compartments does not depend primarily on canonical autophagosome formation under the conditions tested (Figure 5). Following SAR405 treatment, there was an accumulation of LC3-I but no notable reduction in LC3-II abundance, which may reflect the opposing effect of influenza infection on autophagic flux, as PR8 infection, and M2 protein in particular, has been reported to promote LC3-II accumulation by impairing autophagosome-to-lysosome fusion (*68*). In addition, although Vps34 inhibition alone may not fully eliminate autophagy-related PI3P signaling (*101*), the S1-LC3B reporter system, together with the broader PI3K inhibitor wortmannin, provided complementary support for our conclusion. These data do not exclude contributions from other autophagy-related pathways, including microautophagy, chaperone-mediated autophagy (CMA) (*102*, *103*), or non-canonical conjugation of ATG8 to single membranes (*104*). Notably, analysis with KFERQ Finder (*105*) identified four putative CMA-targeting KFERQ-like motifs within the influenza PR8 neuraminidase sequence (fig. S5C), raising the possibility that selective autophagy-related mechanisms could contribute to delivery or processing of neuraminidase-derived intermediates for MHC-II presentation. Together, these findings underscore that non-classical pathways involved in MHC-II–restricted epitope processing remain incompletely understood and underappreciated. Additional studies using 1D6 are underway to interrogate the site of NA25 loading and the mechanism of peptide transport during infection.

Overall, the 1D6 TCR-like antibody provides a new avenue for studying and identifying the cellular machinery that underlies endogenous antigen processing during viral infection, which in the case of influenza, is the primary driver of T_CD4+_ responses (*10*). Specifically, the direct quantification of NA25:A^b^ presentation by 1D6 in both cell-based assays as well as imaging-based localization studies enables approaches that have previously been difficult to achieve with T cell activation assays alone. A key future direction is the combination of 1D6-based fluorescence-activated cell sorting (FACS) with whole-genome CRISPR-Cas9 screening to define the transporters, proteases, trafficking regulators, and MHC-II loading factors that facilitate and regulate endogenous MHC-II presentation. More broadly, this platform provides a generalizable framework to generate TCR-like antibodies and related staining reagents for additional non-canonical MHC-II-restricted epitope, expanding the tools available to directly track naturally processed pMHC-II complexes. These tools could reveal underappreciated pathways by which intracellular viral proteins are processed and displayed to T_CD4+_, with broader implications for understanding antiviral immunity and designing vaccines that more effectively engage helper T cell responses.

## MATERIALS AND METHODS

### Study Design

This study was conducted to generate and characterize a TCR-like antibody to study endogenous MHC-II antigen processing and presentation during influenza A virus infection. Using mRNA-based mouse immunization and pMHC-II tetramer sorting, we generated 1D6, a monoclonal antibody that recognizes the PR8 neuraminidase-derived NA25 epitope presented by I-A^b^. We assessed the specificity and sensitivity of 1D6 using recombinant protein, peptide-pulsed and influenza-infected antigen-presenting cells by ELISA, flow cytometry, surface plasmon resonance, and compared with T_CD4+_ hybridoma activation assays. The structure of 1D6 Fab in complex with NA25:A^b^ was determined by cryo-EM. We then used 1D6 to examine the requirements for endogenous NA25 generation and MHC-II loading by comparing NA25:A^b^ detection after live or inactivated PR8 infection, proteasome inhibition, and GILT deficiency. We also used intracellular 1D6 staining together with subcellular markers to localize NA25:A^b^ complexes and tested whether macroautophagy contributed to NA25 generation using two pharmacologic inhibitors. Experiments that were independently replicated, as well as samples sizes, are noted in the figure legends.

### Mice

WT C57Bl/6 (strain 000664) and BALB/c mice (strain 000651) were purchased from the Jackson Laboratory. Mice were maintained in a specific pathogen-free facility at the Children’s Hospital of Philadelphia (CHOP). Animal protocols were approved by the Institutional Animal Care and Use Committees (IACUC) at CHOP.

### Cell lines and primary APCs

The B6-CIITA and B6-CIITA-E^d^ cell lines were derived from an in-house C57Bl/6 (B6) skin fibroblast cell line by gamma retroviral transduction with human class II transactivator (*Ciita*), promoting endogenous I-A^b^ expression, plus BALB/c-derived *I-E^d^*(B6-CIITA-E^d^ cells only; described previously (*106*)). These cell lines were maintained in Dulbecco’s modified eagle medium (DMEM) containing 5% (D5) or 10% (D10) fetal bovine serum (FBS), 2 mM L-glutamine, and 1x penicillin/streptomycin.

The generation and characterization of T_CD4+_ hybridomas [NA_97-111_/I-A^b^ (‘NA25’ (ref. (*10*))), HA_91-107_/I-A^b^ (‘HA16’ (ref. (*10*))), HA_107-119_/I-E^d^ (‘S1’ (ref. (*20*)))] were previously described. Briefly, antigen-specific T cells were fused with the partner cell line, BWZ.36/CD8α, expressing a nuclear factor of activated T cells (NFAT)-inducible *lacZ* reporter gene. T_CD4+_ hybridomas were maintained in Roswell Park Memorial Institute (RPMI) medium containing 10% FBS, 2 mM L-glutamine, 1x penicillin and streptomycin and 50 µM 2-mercaptoethanol (R10).

Mouse single-B cell culture included feeder cells (NB-21) maintenance and B cell culture as described previously (*35*, *36*). NB-21 cells (clone 2D9) were grown in DMEM supplemented with 10% FCS, 1% penicillin/streptomycin, 1% MEM Non-Essential Amino Acids (NEAA). B cell media (BCM) contained RPMI-1640 supplemented with 10% FBS, 55 µM 2-ME, 1% penicillin/streptomycin, 10 mM HEPES, 1 mM sodium pyruvate, and 1% MEM NEAA.

Immature bone marrow-derived dendritic cells (BMDCs) were generated as previously described (*107*) with the following modifications: media changes were performed every 3 days and BMDCs were harvested on day 7. BMDCs were maintained in R10 supplemented with granulocyte-macrophage colony-stimulating factor (‘GM-CSF’, Shenandoah) at a final concentration of 30 ng/mL. All cells were grown at 37 °C in a 5% CO_2_ incubator.

### Viruses

The PR8 virus was an infectious molecular clone derived from a set of eight ambisense plasmids in the pDZ vector (*108*), which encode each of the PR8 gene segments and are transcribed into both negative sense genomic RNA positive sense mRNA when transfected into cells. WT PR8 virus stock and pDZ plasmids were a kind gift from Scott E. Hensley (University of Pennsylvania). Expansion and titration of the viruses was previously described (*109*). Influenza A/X-31 (H3N2) was a kind gift from Shaon Sengupta (Children’s Hospital of Philadelphia).

Viruses were inactivated using β-propiolactone (*110*) as previously described (*10*).

### mRNA synthesis

Synthetic mRNA was produced by *in vitro* transcription as follows. Plasmid templates encoded either the I-A^b^ α chain or the β chain covalently linked to the NA25 epitope via a flexible Gly/Ser linker, with the design: I-A^b^β signal peptide (27 a.a.)–NA25 epitope (15 a.a.)–(GGGGS)x4–mature I-A^b^ β protein sequence (238 a.a.). The plasmids were each linearized (*111*) and used for *in vitro* transcription (MEGAscript T7 Transcription Kit, Thermo AMB13345) incorporating N1-methylpseudouridine (TriLink N-1081) in place of uridine, and 5’ cap1 structure using CleanCap Reagent AG (3’ OMe) (TriLink N-7413). The mRNA was purified of double-stranded RNA contaminants by adsorption to cellulose (*112*) (Sigma 11363). The mRNA length and integrity were confirmed by agarose gel electrophoresis.

### mRNA immunization and serum analysis

We encapsulated mRNA using the commercial reagent, in vivo-JetRNA+ (Sartorius 101000122, formerly Polyplus), using the manufacturer’s protocol, which forms permanently cationic mRNA liposomes. Three BALB/c mice were immunized with 11 ug I-A^b^ α chain mRNA and 11 µg of NA25-linked I-A^b^ β chain, co-encapsulated with in vivo-JetRNA+, on days 0, 22, and 145. Serum was collected on day 32 for analysis of NA25: A^b^ reactivity. To screen sera, non-NA25-dependent A^b^ reactivity was removed from each serum by first incubating 5 µL serum, diluted 1:100 with 0.1% BSA in PBS, with 30 million A^b^-expressing B6-CIITA fibroblast cells, rotating for 1 hr at 4°C. Adsorbed supernatant was collected after pelleting the cells. Parental B6 fibroblasts not expressing A^b^ were used as an irrelevant adsorption control. Then, 50 µL of each adsorbed serum was used to stain PR8-infected B6-CIITA-E^d^ cells (also A^b+^) for 40 minutes at 4°C, followed by two washes and staining with secondary anti-mouse IgG PE (Jackson 115-116-071). Mice were euthanized on day 151 and spleens were collected for NA25:A^b^ tetramer staining and B cell sorting.

### B cell sort and screening

Splenocytes from immunized mice (see above) were DNase I-treated and first stained with 400 uL of PBS with 2% BSA with 2 mM EDTA containing anti-CD16/CD32 (20 µg/mL) and a decoy pMHC-II tetramer, NP_311-325_:A^b^ (20 µg/mL) for 20 m at 4°C. Then, an equal volume of buffer containing NA25:A^b^ -PE and NA25:A^b^ -BV421 tetramers was added for a final concentration of 2 µg/mL each. Cells were washed, stained with Live/Dead Aqua, washed, and stained with a cocktail of antibodies each at 2 µg/mL in 800 µL of buffer: B220 BV605 (Biolegend 103241), IgG1 FITC (BD 553443), IgG2b FITC (Biolegend 406706), IgG2a/c FITC (Biolegend 407106), and IgM PerCP-eFluor 710, as well as the following not used for gating: CD3 Alexa Fluor 700 (Biolegend 100216), CD4 BV786 (BD 563727), and a dump channel (Ly6-C, Ly6-G, and F4/80 PE-Cy7). Surface IgG^+^ B cells (Aqua^-^ B220^+^ IgM^-^ IgG1/2a/2b^+^) that were negative for the decoy NP_311-325_:A^b^ tetramer but double positive for the NA25:A^b^ tetramers were singly sorted into flat 96-well plates seeded 1 day before with 2,000 NB-21 cells per well in BCM and supplemented several hours before with mouse IL-4 at a final concentration of 2 ng/mL in a final volume of 200 µL BCM. B cell/NB-21 co-cultures were cultured at 37°C 5% CO_2_ for 10 days, with 50% medium replacement on day 2 and 100% medium replacement on days 3-8. On day 10, the supernatant was collected and stored at 4°C, and the cells were left in the plates and frozen at -80°C for later B cell receptor cloning.

B cell/NB-21 supernatants were screened for NA25:A^b^ reactivity by a combination of ELISA and flow cytometry. For ELISA, Nunc MaxiSorp plates (Thermo 439454) were coated with 2 µg/mL streptavidin overnight at 4°C, washed with wash buffer (0.05% Tween-20 in PBS) and blocked with 2% BSA (Sigma A7030) in PBS. Then, 0.4 µg/mL of NA25:A^b^ biotinylated monomer (NIH Tetramer Core Facility) was added to the wells for 1 hr, shaking, at room temperature. After washing, supernatants were added at a ≤ 1:12 dilution in 2% BSA buffer and incubated 1 hr, shaking, at room temperature. After washing, 0.16 µg/mL donkey anti mouse whole IgG HRP (Jackson 715-035-150) was used as a secondary antibody, and TMB was used to develop the signal. In another version of the ELISA, 10 µg/mL synthetic peptide was used as the capture antigen in place of streptavidin and biotinylated pMHC-II.

For flow cytometry screening, B6-CIITA-E^d^ fibroblasts were infected with IAV PR8 at 25 HAU per million cells then cultured in suspension, rotating for ∼18 hours. Cell suspensions were passed through a 100 µm filter and plated at 0.5 million cells per well in a U-bottom 96-well plate. Cells were first stained with Live/Dead Aqua, then a 1:2 dilution of B cell/NB-21 supernatants for 40 m, then washed and stained with 2 ug/mL secondary goat anti-mouse Ig Biotin (Jackson 115-066-068) for 30 m, and then washed and stained with 2 µg/mL streptavidin-PE (Jackson 016-110-084) for 20 m; all stains were at 4°C. %IgG^+^ (PE^+^) events were quantified within live (Aqua^-^) singlets.

### Antibody cloning and expression

The B cell receptor variable region was amplified and cloned from B cell clone 1D6 using a protocol described by von Boehmer et al. (*113*) for single B cells, with the following modifications. To account for our starting material being a frozen cell culture of approximately 50 µL, the B cell/NB-21 cell plate was thawed, and Lysis Buffer and RT Mix I were immediately added to reach a total volume of ∼140 µL per well. This lysate was diluted 1:10 in the same buffer, and 11 µL of the dilution was combined with RT Mix II to proceed with the RT and PCR reactions. The heavy and light chain variable regions of B cell clone 1D6 were cloned into mouse heavy and light chain expression vectors used previously (*113*) by SLIC cloning and then sequenced.

Recombinant antibody clone 1D6 was expressed by co-transfecting ExpiCHO-S cells with equal amounts of heavy and light chain plasmids using the ExpiFectamine CHO Transfection Kit per the manufacturer’s protocol and was purified by Protein G chromatography.

### Synthetic peptides

The following synthetic peptides were used: NA_97-111_ (GDVFVIREPFISCSH)(*10*), HA_91-107_ (RSWSYIVETPNSENGIC)(*10*), and NP_276-292_ (LPACVYGPAVASGYDFE)(*10*). All peptides were obtained lyophilized at > 85% purity from GenScript and dissolved in DMSO at a stock concentration of 10 mg/mL or 20 mg/mL. Peptide pulsing was performed by incubating cells with 20 µg/mL peptide overnight or 300 µg/mL peptide for 1 hr at 37°C, followed by three washes and then downstream staining.

### *In vitro* antigen-presentation assays

For *in vitro* APC infection, cells were infected with indicated influenza viruses at different MOIs as specified in 0.1%BSA in PBS without Ca/Mg (0.1% BSA/PBS). Cells were incubated with viruses at 37°C for 45 minutes to allow for viral attachment and then changed to complete medium for overnight (16-20 hours) infection, followed by harvesting for flow staining or imaging.

For proteasome inhibition assays, APCs were pre-treated with marizomib (MCE HY-10985) at indicated concentrations for 15 minutes at 37°C and washed twice before infection with virus at 5 HAU per 1 million cells as described above. APCs were further incubated with T_CD4+_ hybridomas and co-cultured overnight or changed to complete medium with 1M HEPES pH7.4 (Gibco) and rotated at 37°C end-over-end for 16-20 hours before collection for flow staining.

For macroautophagy inhibition assays, APCs were pre-treated with SAR405 (MCE HY-12481) or wortmannin (MCE HY-10197) at the indicated concentrations in 0.1% BSA/PBS for 1 hr at 37°C prior to influenza virus infection. APCs were then infected with influenza virus at the indicated MOIs as described above. SAR405 or wortmannin was maintained throughout viral attachment and overnight infection.

For transient expression of S1-LC3 and S1-LC3(G120A), APCs were transfected with plasmid DNA using Lipofectamine 2000 (ThermoFisher) according to the manufacturer’s recommendations. Cells were incubated at 37°C for 20-24 hours, pelleted, and washed once with ice-cold 1x PBS. Cells were then incubated with ice-cold 0.1M glycine (pH 2.5) on ice for 30 sec, followed by immediate neutralization with 1/10 volume of 1M Tris (pH 8), and 10x volume of ice-cold complete medium. Cells were washed again with complete media and resuspended in ice-cold D5 with HEPES, with SAR405 at the indicated concentrations or DMSO as a control. Cells were treated with SAR405 overnight with end-to-end rotation. The following day, cells were harvested, fixed with 0.05% PFA on ice for 15 minutes, and washed extensively. Fixed APCs were then incubated with T_CD4+_ hybridomas for 6 hours before plate reading.

### Preparation of mouse tissues

Mouse bone marrow was collected by flushing it from the femur and tibia with R10 medium. Cell suspension was passed through a 40 µm filter followed by ACK lysis. BMDCs were plated by plating 2 million cells in 10 mL R10 medium in 10 cm tissue culture plates with GM-CSF.

Spleens were collected in R10 and manually homogenized through a 70 µm cell strainer using the hard end of a 1-mL syringe plunger and subsequently washed with 10 mL of FACS buffer (PBS + 1% FBS + 2 mM EDTA). Homogenates were then spun down at 4 °C and resuspended in 3 mL of ACK lysis buffer, left at RT for 3 minutes, and topped up with cold PBS before spinning again. Cell pellets were resuspended in 10 mL of FACS buffer and passed through a 40 µm strainer to prepare a single cell suspension.

All primary cell samples were maintained on ice during preparation (except as otherwise specified) and until analysis.

### Immunofluorescence microscopy cell preparation

To prepare coverslips for immunofluorescent imaging, autoclaved coverslips were placed into 24-well dishes and coated with 50 μg/mL collagen in 0.02 M acetic acid (B6-CIITA cells) or 0.5 mg/mL poly-L-lysine in 1X PBS without calcium/magnesium (B6-CIITA-E^d^ cells) for 1 hour at room temperature. B6-CIITA (no E^d^) cells also stably express an ER-resident mNeonGreen (*114*). During coverslip coating, B6-CIITA (1D6 + LAMP-1/GM130) or B6-CIITA-E^d^ (1D6 + Y-Ae + LAMP-1) cells were infected with IAV strain PR8 or strain X31 at a concentration of 5 HAU per 1 x 10^6^ cells for 45 minutes at 37°C, gently resuspending cells throughout viral attachment. After viral attachment, cells were washed to remove free virions and plated onto coated coverslips in D5 media for 16 hours at 37°C. The following day, coverslips were washed twice with 1X PBS with calcium/magnesium and fixed using BD Cytofix/Cytoperm Fixation/Permeabilization Kit (BD Biosciences Cat#554714) for 20 minutes at 4°C. Cells were washed three times with 1X BD Perm/Wash buffer (BD Biosciences Cat#554723) and stained with anti-LAMP-1 (Cell Signaling Technologies, clone E5N9Z, 1:200) or anti-GM130 (Cell Signaling Technologies, clone E9Z6S, 1:400) and AF647-conjugated 1D6 (2 μg/mL) diluted in 1X BD Perm/Wash buffer for 1 hour at room temperature. After the primary stain, cells were washed three times and stained with secondary antibody comprising AF790-conjugated anti-rabbit IgG (ThermoFisher, Cat#A11369, 1:500) (1D6 + LAMP-1/GM130) or Coralite Plus 488-conjugated anti-rabbit IgG (Proteintech, Cat#RGAR002, 1:500) (1D6 + Y-Ae + LAMP-1) and Coralite Plus 647-conjugated anti-mouse IgG (Proteintech, Cat#RGAM005, 1:500) diluted in 1X BD Perm/Wash buffer for 1 hour at room temperature. After the secondary stain, cells were washed three times and stained with tertiary antibody comprising AF568-conjugated anti-PR8 neuraminidase (Clone NA2-1C1, 5 μg/mL) and AF790-conjugated anti-Eα_52-68_:A^b^ (ThermoFisher, Cat#14-5741-82, 2 μg/mL) (1D6 + Y-Ae + LAMP-1 only) diluted in 1X BD Perm/Wash buffer for 30 minutes at room temperature. After the tertiary stain, cells were washed twice with 1X BD Perm/Wash buffer and once with 1X PBS without calcium/magnesium. Cells were then stained with DAPI (Abcam, Cat#AB229549, 1:1000) diluted in 1X PBS without calcium/magnesium for 5 minutes at room temperature prior to a final three washes and mounting using ProLong Diamond Antifade Mountant (Invitrogen, Cat#P36970). All cells were imaged on a Leica Stellaris 5 confocal microscope in the University of Pennsylvania Perelman School of Medicine Cell and Developmental Biology Microscopy Core Facility (RRID:SCR_022373).

### Immunofluorescence microscopy analysis

The Fiji distribution of ImageJ2 (https://imagej.net/software/fiji/) was used for all image analyses. To quantify colocalization of 1D6, LAMP-1, GM130, and Y-Ae signals, the PR8 neuraminidase channel was used to manually draw cell outlines, adding each cell as an individual Region of Interest (ROI). Afterwards, 100 pixel sliding parabola background corrections were performed for each of the 1D6, LAMP-1, GM130, and Y-Ae channels. Coloc2 was then used to quantify the Pearson correlation and M1 Mander’s coefficient.

### Hybridoma activation assay

For viral infection, APCs were collected and resuspended at 1 x 10^6^ cells/mL in 0.1%BSA/1x PBS with influenza viruses at indicated MOIs and incubated at 37°C for 45 minutes to allow for viral attachment. Cells were pipetted every 15 minutes during this incubation. APCs were then washed twice and resuspended at 1 x 10^6^ cells/mL in R10 media. For untreated control or peptide-treated conditions, APCs were collected and resuspended at 1 x 10^6^ cells/mL in R10 media directly. T_CD4+_ hybridomas were harvested and resuspended at 1 x 10^6^ cells/mL in R10 media and left on ice till plating. To a 96-well black flat-bottom microplate (Corning 3916), 50 µl of APCs were plated into respective wells. For infected or untreated cells, extra 50 µl of R10 were added. For peptide-treated conditions, 50 µl of 40 µg/mL synthetic peptide diluted in R10 were added to each corresponding well to a final concentration at 10 µg/mL. 100 µl T_CD4+_ hybridomas were then added into each well and co-cultures were incubated at 37°C for 16-20 hours. After incubation, co-cultures were lysed with a substrate buffer containing 1.25% Triton X-100, 22 µg/mL 4-methyl-umbelliferyl-β-D-galactopyranoside (Sigma-Aldrich), 38.5 µM 2-mercaptoethanol and 9 mM MgCl_2_ in PBS for 3 hours at 37°C. After incubation, fluorescence was quantified at 365/445 nm using a microplate reader. Hybridoma assays consisted of at least three technical replicates and were performed several independent times where indicated.

### Antibody/hybridoma competition assay

B6-CIITA-E^d^ fibroblasts were infected with influenza PR8 at an MOI of 10 as described above. At 16-18 hours post infection, cells were harvested and plated into black, flat-bottom 96-well microplate as described above. Unconjugated 1D6 or isotype control antibody was serially diluted and added to wells containing APCs at 2x the desired concentration. Cells were incubated with antibody at 37°C for 2 hours to allow for blocking. After the initial blocking period, T_CD4+_ hybridomas were added as described above and incubated with APCs in the continued presence of antibody for an additional 16-20 hours before reading out hybridoma activation as above.

### Peptide:MHC-II monomers and tetramers

The following pMHC-II tetramers and monomers were obtained from the NIH Tetramer Core Facility: NA25:A^b^ PE tetramer, NA25:A^b^ BV421 tetramer, NA25:A^b^ biotin monomer, and NP_311-325_:A^b^ APC tetramer.

For in-house generation of linked NA25:A^b^ monomer, a codon-optimized DNA sequence encoding the NA25 peptide fused to the mouse A^b^β chain was synthesized by Twist Bioscience. The construct contained the endogenous *H2-Ab1* signal peptide followed by the NA25 peptide linked to the mature mouse A^b^β chain (H2-Ab1, Uniprot P14483). This was followed by a P2A peptide sequence and a second open reading frame encoding the H2-Aa signal peptide fused to the mature mouse A^b^α chain sequence (H2-Aa, Uniprot P14434). The transmembrane domains were removed to generate soluble A^b^. Jun and Fos leucine zipper sequences were appended separately to the C-terminal of the α and β chains to promote heterodimer formation. An 8x-His tag was fused to the C-terminal of α chain to facilitate purification.

Expi293F cells (ThermoFisher Scientific) were cultured, maintained in Expi293 Expression Medium (Gibco), and transiently transfected according to the manufacturer’s recommendations, shaking at 125 rpm with 8% CO_2_. Briefly, healthy cells (≥95% viable) at 3.0 x 10^6^ cells/mL were transfected with ExpiFectamine 293 and the appropriate construct DNA at a final concentration of 1 µg DNA per mL of culture and maintained at 37°C. Transfection kit enhancers 1 and 2 were added 18 h post-transfection at the recommended volumes. Cultures were harvested after 5 days of transient transfection via centrifugation at 3000 x g for 30 minutes at 4°C, after which the supernatant was filtered through a 0.22 µm polyethersulfone membrane (Millipore).

To prepare EDTA-compatible Ni-IMAC resin (Pierce) for His-tag protein capture, slurry was buffer exchanged to equilibration buffer (50 mM monosodium phosphate, 300 mM sodium chloride, 10 mM imidazole, pH 8.0). The equilibration buffer was removed and harvested Expi293 supernatant was incubated with the washed beads. The cell supernatant was rotated end-over-end with beads for 1 hour at 4°C. After incubation, the supernatant was run through a disposable gravity flow column (Bio-Rad) and rinsed once with 1 column volume (CV) of equilibration buffer. The settled resin was then washed with 3 CV of wash buffer (50 mM monosodium phosphate, 300 mM sodium chloride, 20 mM imidazole, pH 8.0). The bound protein was eluted with 3 x 3 mL of elution buffer (50 mM monosodium phosphate, 300 mM sodium chloride, 500 mM imidazole, pH 8.0). Elutions were pooled and dialyzed overnight using a 10 kDa dialysis cassette (ThermoFisher) in 1x PBS at 4°C. The following day, dialyzed protein was further concentrated using a 10 kDa centrifugal filter (Amicon).

### SDS-PAGE and western blot

Cells were collected and lysed in 1x RIPA buffer (1% Triton X-100, 100 mM HEPES, 1 mM CaCl_2_) and incubated at 4°C on a rocker for 30-60 minutes. Following incubation, lysates were clarified through centrifugation at 12000 rpm x 10 minutes at 4°C. Lysate samples were then denatured and reduced through the addition of 4X LDS Sample Buffer (NuPAGE) and 10X Sample Reducing Agent (NuPAGE), respectively. Samples were denatured at 90-95°C for 10 minutes before loading in 12% Bis-Tris gels (NuPAGE) (with MES buffer) or 4-12% Bis-Tris gels (NuPAGE) (with MES buffer). Gels were run at 150 V. When complete, gels were stained in Coomassie protein stain (Abcam) directly for SDS-PAGE.

For western blot, protein was transferred to nitrocellulose membranes using a Trans-Blot Turbo Transfer System (Bio-Rad) set at 25 V, 1.0 A, 30-40 minutes, with a constant voltage. Following transfer, membranes were incubated in Intercept (TBS) Blocking Buffer (LICORbio) for 60 minutes at room temperature. Blocking buffer was then removed and replaced with primary antibody diluted in Intercept T20 Antibody Diluent (LICORbio). Membranes were incubated with primary antibody (anti-LC3B antibody: Sigma-Aldrich L7543, working concentration 2 µg/mL; anti-β-actin antibody: Santa Cruz sc-47778, working concentration 2 µg/mL) overnight at 4°C, with gentle rocking. The following day, membranes were washed three times with TBS-T (TBS + 0.05% Tween 20) and incubated with secondary antibodies (IRDye 800CW Donkey anti-Mouse IgG Secondary Antibody: LICORbio 926-32212; IRDye 680RD Goat anti-Rabbit IgG Secondary Antibody: LICORbio 926-687071) diluted in Intercept T20 Antibody Diluent for 45 minutes at room temperature. After three washes with TBS-T, membranes were imaged using the LiCor Odyssey or Azure 500 (Azure Biosystems) fluorescent imager.

### Surface plasmon resonance

SPR experiments were performed in duplicate using a Biacore T200 instrument (Cytiva) in SPR buffer [0.01M HEPES (pH7.4), 0.15M NaCl, 3 mM EDTA, 0.05% (v/v) Surfactant P20; Cytiva HBS-EP BR100826]. Approximately 50-200 resonance units (RU) of biotinylated NA25:A^b^ or NP47:A^b^ as a specificity control, was immobilized on a streptavidin-coated chip (XanTec Bioanalytics, SCBS SAHC200M) at a flow rate of 10 µL/min. Serial dilutions of 1D6 antibody, ranging from 0 to 100 nM, were injected over the chip surface at 25°C at a flow rate of 30 µL/min for 180s, followed by a 900s dissociation phase. The surface was regenerated with 1M MgCl_2_ for 100s at a flow rate of 25 µL/min, followed by a 60s stabilization period. SPR sensorgrams and equilibrium dissociation constants (K_D_) were analyzed using the surface-bound analysis settings in the Biacore T200 Evaluation Software version 3.2.1 (Cytiva). Representative SPR sensorgrams were prepared using GraphPad Prism.

### Cryo-EM sample preparation, data collection and data processing

1D6 Fab was generated by digesting 1D6 IgG with papain (Sigma). The Fab fraction was further purified by incubating with Protein A resin (Cytiva). Purified NA25:A^b^ was incubated with 1D6 Fab at a molar ratio of 1:4 on ice for 2 hours followed by Size Exclusion Chromatography via a Superdex 200 Increase 10/300 GL column. Corresponding fractions were concentrated to 0.5 mg/mL. 4 µL of sample was applied to glow-discharged UltrAufoil R1.2/1.3 grid and blotted for 4s followed by plunge-freezing into liquid-nitrogen-cooled ethane. Clipped grids were imaged on a Thermo Scientific Glacios (200kV) equipped with a Falcon 4 detector. Data processing was performed in CryoSARC (*115*). 2896 movies were motion corrected followed by CTF (Contrast Transfer Function) estimation. Blob picked particles were cleaned by several rounds of 2D classification and 3D classification. 77,198 particles were used in local refinement. Resolution was 4.5Å by the FSC 0.143 criterion and 5.1Å by the FSC 0.5 criterion. The final locally refined density map was lowpass-filtered to the appropriate FSC 0.5 resolution of 5.1 Å.

1D6 VH/VL models were generated using SWISS MODEL (*115*). Initial model for NA25:A^b^ was PBD 6MNG. Combined initial model of 1D6/NA25:A^b^ was fitted to the density map. CDR loops and peptide were built in COOT (*116*). Figure panels were generated in ChimeraX (*117*).

### Disulfide bond prediction

Because no experimental structure is available for PR8 NA, PR8 NA was aligned to the structurally characterized A/California/04/2009 H1N1 NA (Uniprot C3W5S3) using BLASTp (*118*). The two proteins showed 80% sequence identity and 88% positives, with strong conservation across the NA25-containing region. The A/California/04/2009 H1N1 NA structure (PDB: 3TI5) was thus used as a homologous structural template. Candidate disulfide bonds were then evaluated using Disulfide by Design 2.0 (*119*), which predicted that the cysteine residues corresponding to PR8 Cys109 and Cys114 are compatible with disulfide bond formation.

### Quantification and statistical analysis

All statistical analyses were performed using GraphPad Prism. Data are presented as mean ± SEM. Comparisons involving two independent variables were performed using two-way analysis of variance (ANOVA) with Šídák’s multiple-comparisons test. For comparisons between two groups, an unpaired two-tailed Student’s *t* test was used. IC_50_ values were calculated using a four-parameter logistic (4PL) sigmoidal model.

For fluorescence colocalization analysis, Pearson’s correlation coefficient was used to assess the linear relationship and signal proportionality between two channels; Manders’ M1 coefficient was used to quantify co-occurrence. For all statistical readouts, \**P* < 0.05, \*\**P* < 0.01, \*\*\**P* < 0.001, and \*\*\*\**P* < 0.0001; ns, not significant.

## Supplementary Materials

Figs. S1 to S5

## Acknowledgments

We thank members of the Eisenlohr lab for their insights and feedback throughout the development of this work; Dr. Amelia Escolano and her lab for helpful discussion regarding B cell isolation and antibody cloning; Dr. Hoang Anh Pham for discussion regarding pMHC-II monomer protein construct design and expression; Dr. Yi Sun for assistance with SPR sample preparation and data analysis. Flow cytometry data and single cell sorting were acquired using the Children’s Hospital of Philadelphia Flow Cytometry Core Facility (RRID: SCR_0097826). We thank the Wistar Institute Molecular Screening and Protein Expression Core Facility (RRID: SCR_024978), with support provided by Cancer Center Support Grant (P30 CA010815), and specifically thank Joel Cassel for providing technical support on SPR data acquisition. Immunofluorescent microscopy analysis was performed at the University of Pennsylvania Cell and Developmental Biology Microscopy Core (RRID: SCR_022373). We thank the NIH Tetramer Core Facility for tetramer and monomer reagents. Cryo-EM data was acquired at The Integrated Structural Biology Shared Resources at Thomas Jefferson University. This work was supported by National Institutes of Health instrument award S10 OD030457 for the acquisition of the Thermo Scientific Glacios/Falcon 4 instrument. This work was supported by National Institutes of Health instrument award S10 OD012063 for the acquisition of the T200 Surface Plasmon Resonance. Model illustrations were created using BioRender.

## Author contributions

Conceptualization: MJH, LCE, LY, SDC

Methodology: MJH, LY, SDC, EH

Investigation: MJH, LY, SDC, EH, JD, MEO, KSK, RN, CAB, NL, CB

Visualization: MJH, LY, SDC, JD

Funding acquisition: LCE, JP

Supervision: MJH, LCE, JP

Writing – original draft: LY, MJH, SDC, LCE

Writing – review & editing: MJH, LCE, LY, SDC, JD, JP

## Data, code, and materials availability

All data needed to evaluate the conclusions in the paper are present in the paper or the Supplementary Materials. All materials used or generated in this study are commercially available or will be supplied upon reasonable request by Penn MTA to L.C.E. Penn MTA templates are available at https://researchservices.upenn.edu/areas-of-service/systems/research-inventory-system/material-transfers/ and outline the key terms and conditions that Penn typically requires for the transfer of its materials to outside parties. No custom scripts or proprietary code was generated for this study.

## Supplementary Table and Figures

**Fig. S1.**
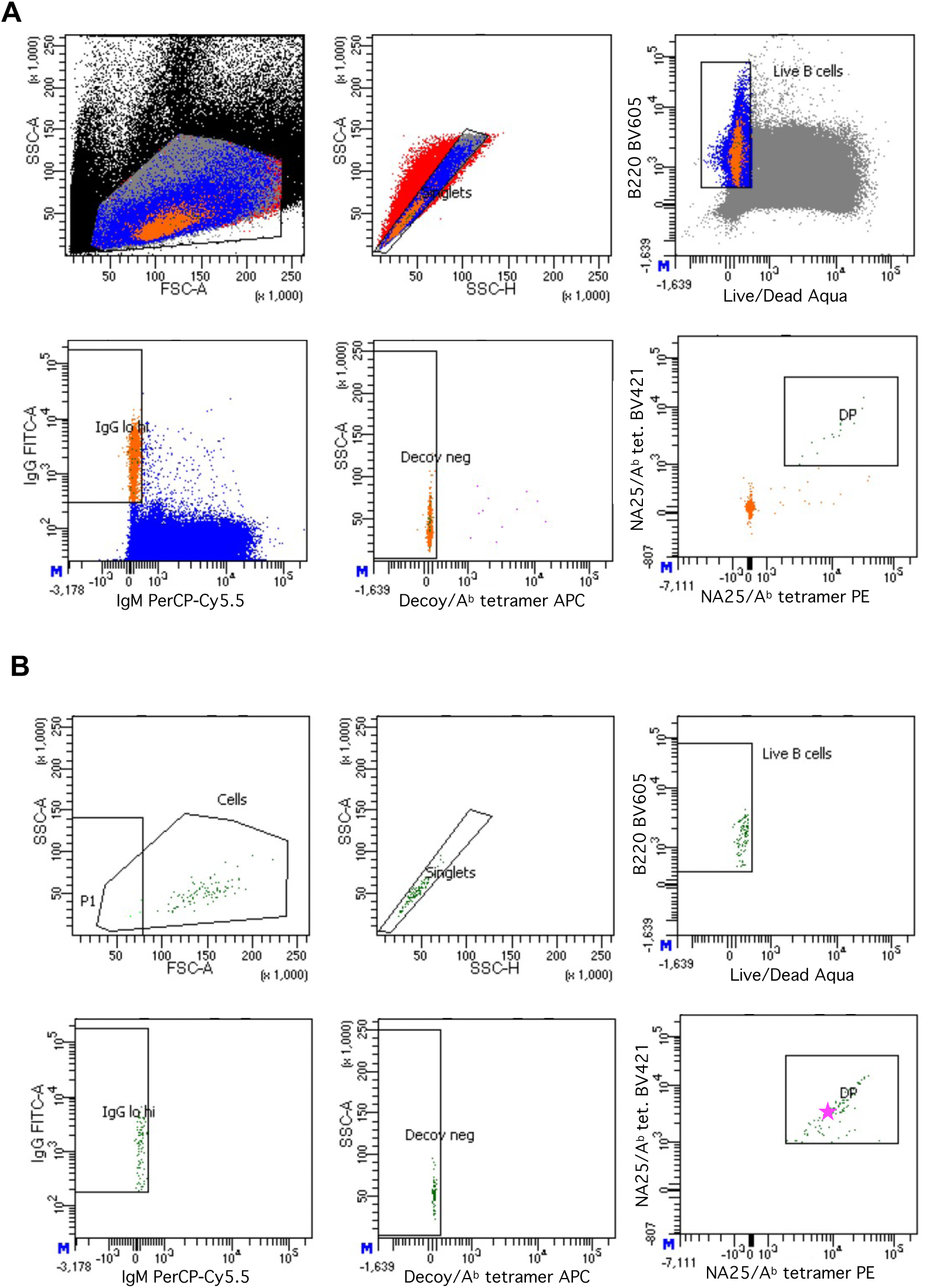
Gating strategy and index data from sort of NA25:A^b^-specific B cells. (**A**) Gating strategy to identify and sort NA25:A^b^-specific B cells from mouse spleen. (**B**) Index sort data of cells singly sorted onto NB21 feeder cell monolayers. The B cell giving rise to the 1D6 antibody is indicated with a magenta star in the bottom right plot. Plots proceed from parent to daughter going left to right, then top to bottom. Decoy=NP_311-325_ IAV epitope. DP=double tetramer positive. P1=irrelevant gate.

**Fig. S2.**
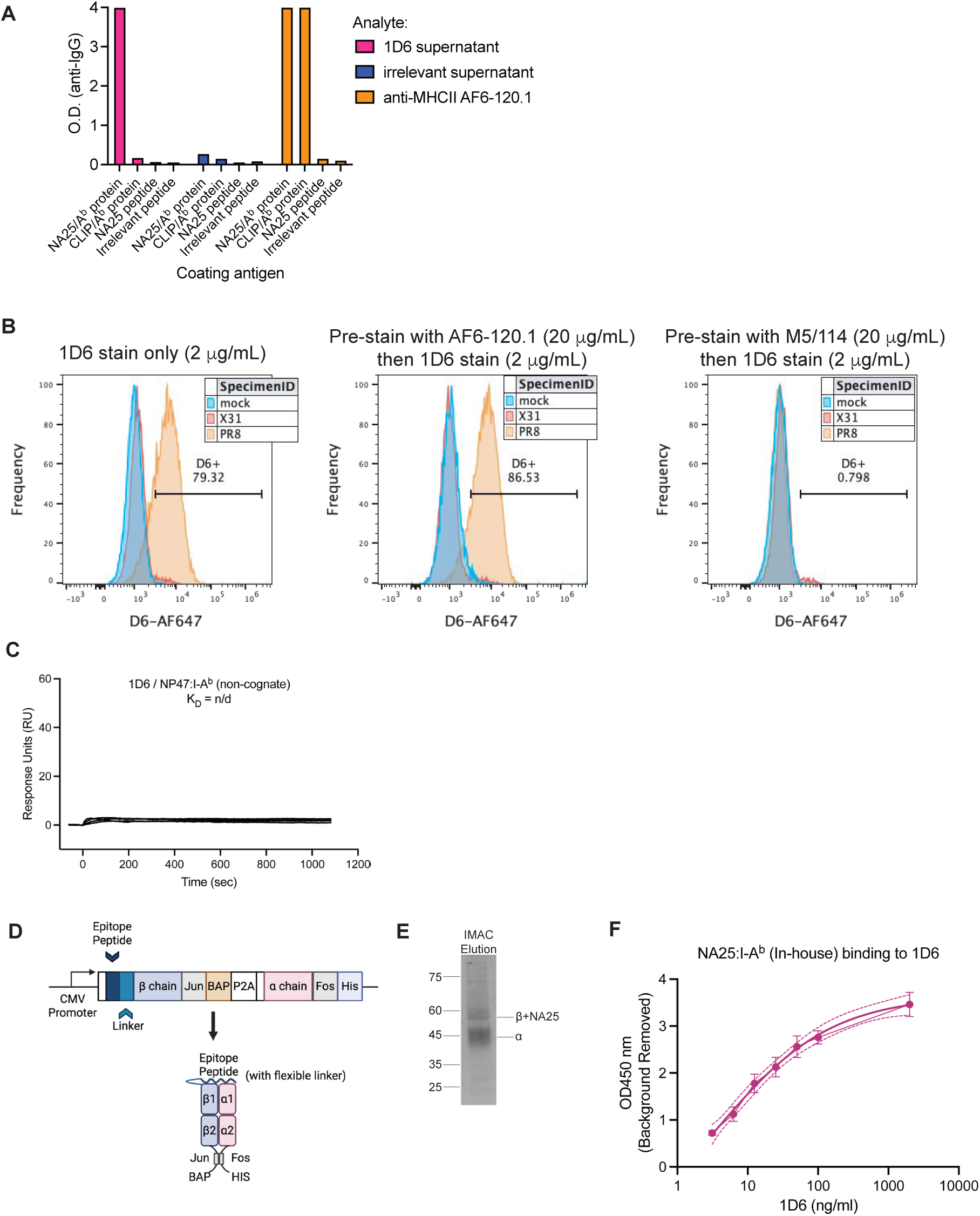
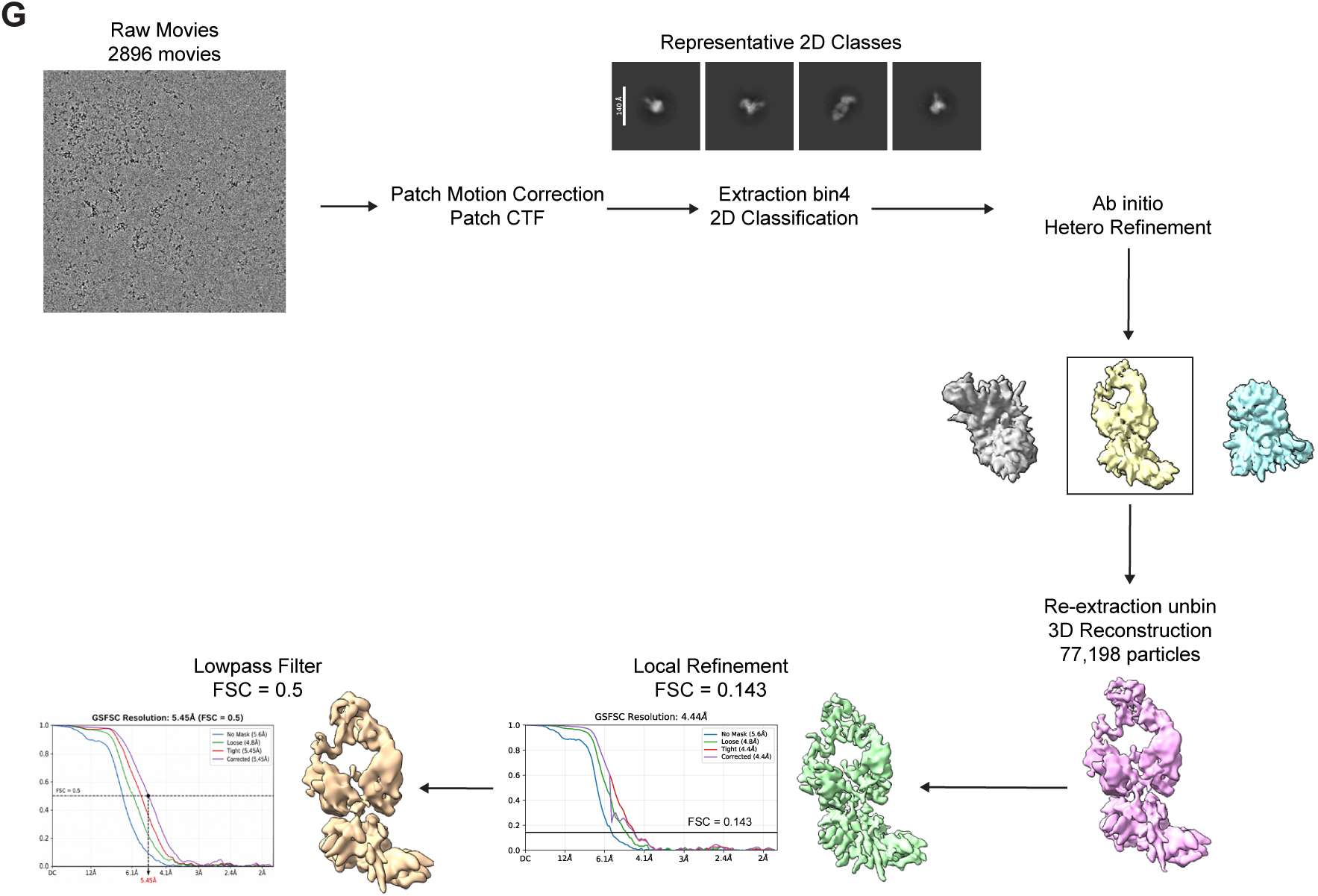
1D6 binds to NA25:A^b^ with high specificity. (**A**) 1D6-containing supernatant from sorted B cell/NB-21 co-culture was tested for specific recognition of Ab protein bound to NA25 but not human CLIP peptide, and lack of binding to soluble NA25 peptide. (**B**) 1D6 recognition of IAV PR8-infected A^b^+ mouse fibroblasts is abrogated by pre-incubation with unlabeled M5/114 but not AF6-120.1 antibody. (**C**) Representative SPR sensorgrams showing no detectable binding of 1D6 to the non-cognate NP47:A^b^ monomer. (**D**) Schematic of the soluble MHC-II DNA construct design used for monomer production. Method and graphic adapted from (*46*). (**E**) SDS-PAGE analysis of eluted, denatured NA25:A^b^ confirming the expected molecular size. (**F**) ELISA showing 1D6 binding to in-house produced soluble NA25:A^b^ monomer. The binding curve was fitted using a sigmoidal four-parameter logistic (4PL) model, from which the EC_50_ was calculated. (**G**) Cryo-EM data processing workflow.

**Fig. S3.**
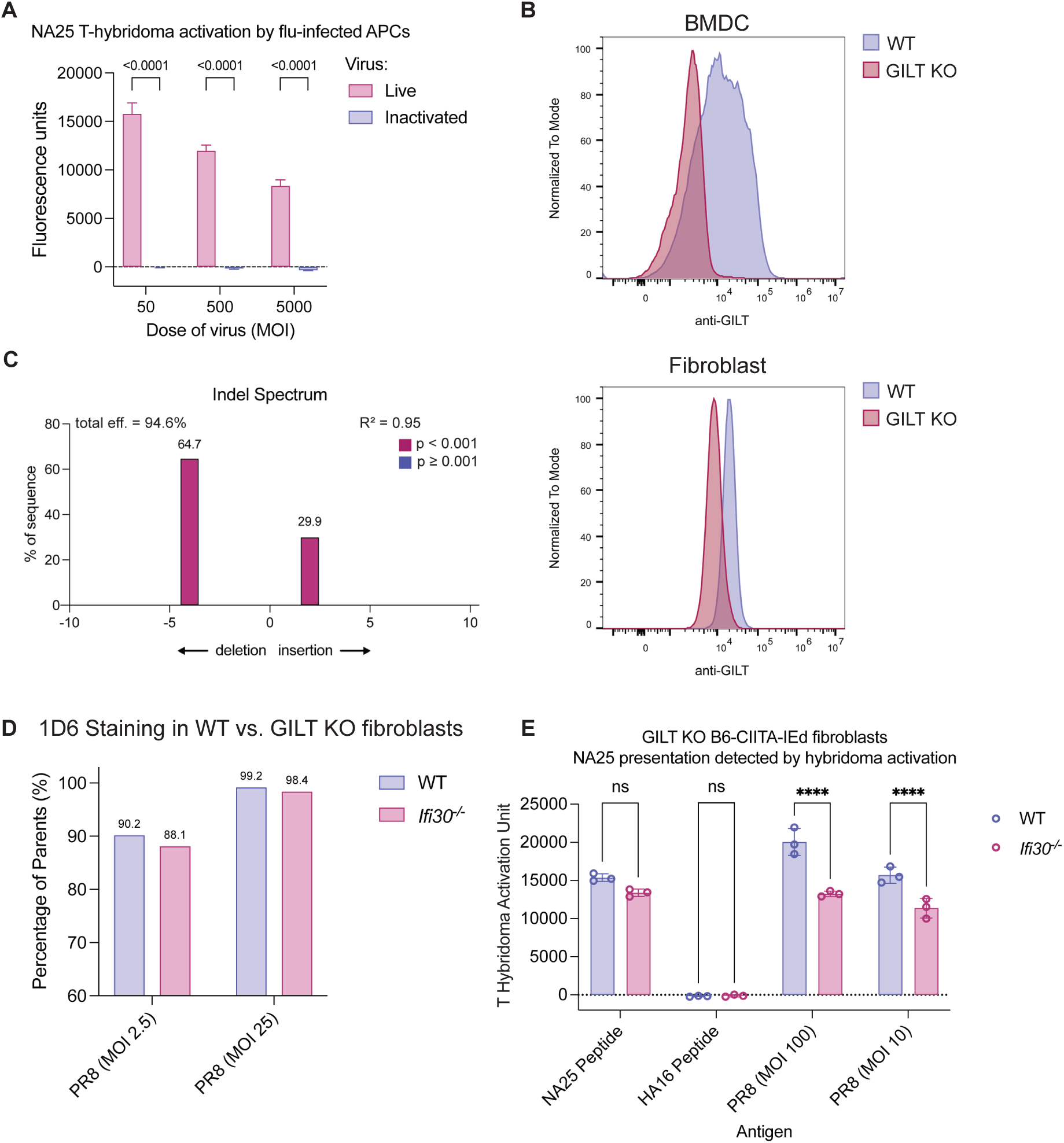
Endogenous NA25 presentation indicated by hybridoma activation, impact of marizomib on NA expression, and validation of GILT knockout. (**A**) Representative data for two independent experiments. B6-CIITA-E^d^ fibroblasts were treated with live or BPL-inactivated PR8 at the indicated MOIs. Data were background subtracted (uninfected control). Bars show the mean +/- s.d. and p-values are shown from a two-way ANOVA with Šídák’s multiple comparison test comparing live and inactive PR8-treated conditions. (**B**) Representative histograms for intracellular GILT staining in GILT KO and WT BMDCs. (**C**) (Left) Graphical representation of Sanger sequencing from selected GILT KO B6-derived fibroblast clone. (Right) Representative histograms for intracellular GILT staining in GILT KO and WT fibroblasts. (**D**) Percent surface 1D6^+^ cells (of total NA^+^ cells) among cells infected with PR8 at indicated MOIs. (**E**) NA25-specific T_CD4+_ hybridoma activation in WT and GILT KO B6-CIITA-E^d^ fibroblasts incubated with indicated peptides or infected with live viruses. Representative of two independent experiments.

**Fig. S4.**
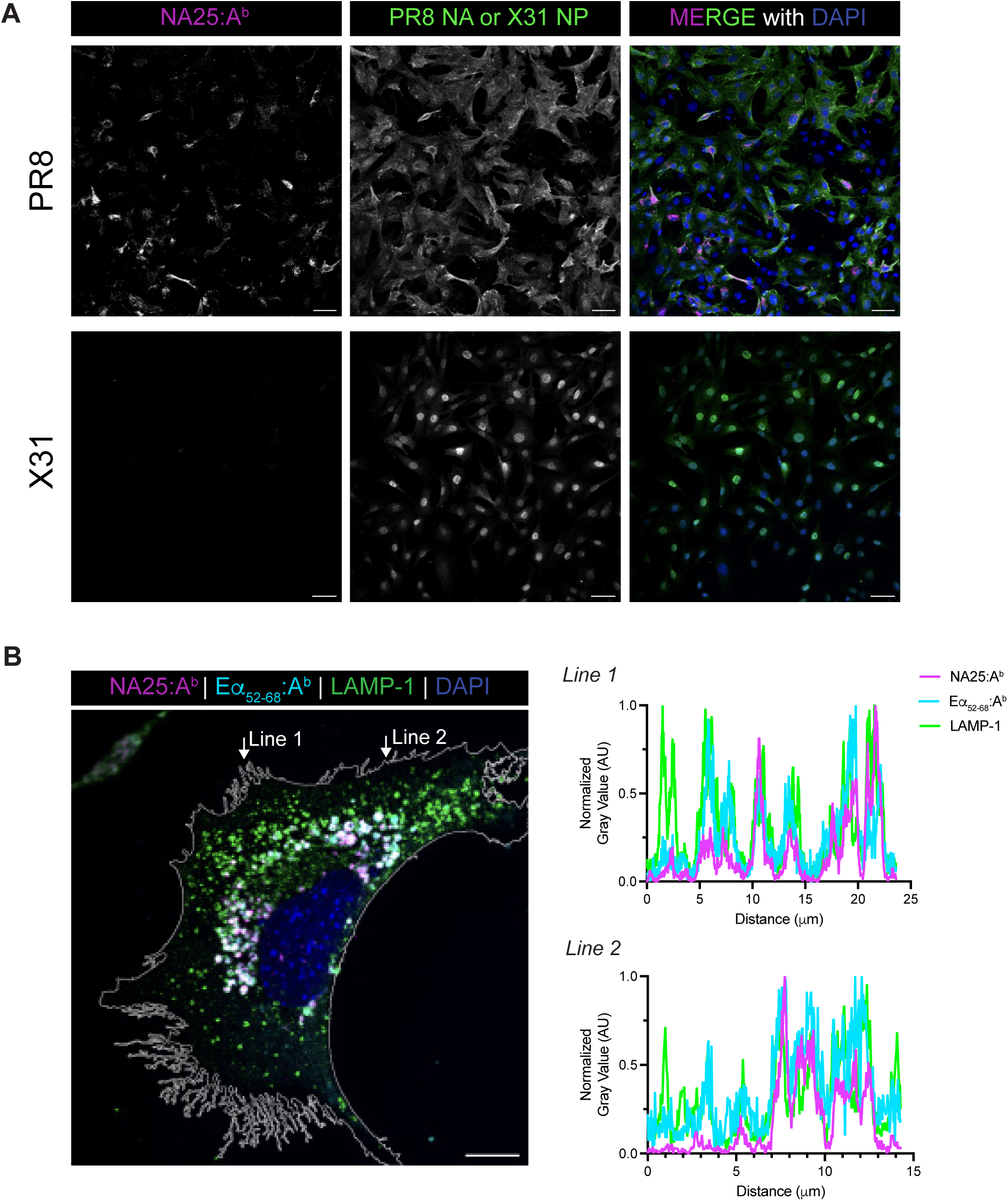
Supporting data for Figure 4. (**A**) Representative composite and single channel immunofluorescent images of NA25:A^b^ complexes (stained with 1D6, magenta) and neuraminidase (IAV PR8, green) or nucleoprotein (IAV X31, green) in B6-CIITA cells, with DAPI used as a counterstain (blue). Scale bar indicates 50 µm. (**B**) Intensity (grey value) line profile of each individual channel beneath the Line 1 and Line 2 arrows in the immunofluorescent image from Figure 4E. Mean grey values along the lines were normalized to the maximal signal in each corresponding channel.

**Fig. S5.**
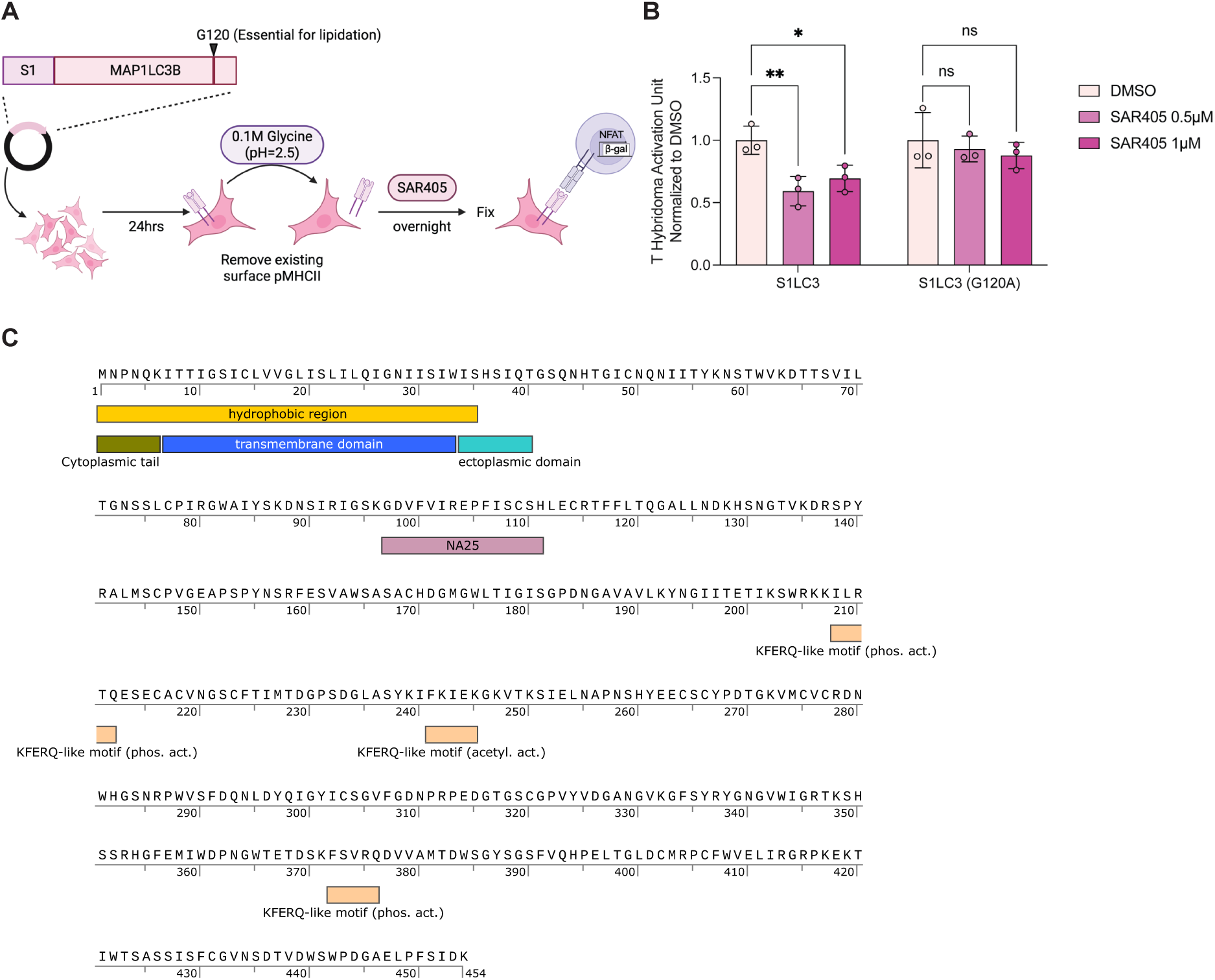
SAR405 activity validation and predicted CMA-targeting motifs in influenza neuraminidase. (**A**) Schematic depicting the S1-LC3B plasmid DNA transfection and SAR405 inhibitory activity quantification via T cell hybridoma assay. (**B**) S1-specific T cell hybridoma activation normalized to the DMSO-treated group. Two-way ANOVA with Šídák’s multiple comparisons test. (**C**) Influenza A/Puerto Rico/8/1934 H1N1 (Mt. Sinai) NA protein sequence with the NA25 epitope and predicted CMA-targeting KFERQ-like motifs labeled. The sequence shown here corresponds to the Mount Sinai strain and differs from the Uniprot P03468 sequence by a single amino acid substitution (I8T), resulting in the loss of one putative KFERQ-like motif in the transmembrane domain.

**Table S1.** Cryo-EM data collection statistics.

| **Data Collection and Processing** | **1D6 Fab/NA25:A<sup>b</sup> complex** |
| --- | --- |
| **Electron Microscope** | Glacios |
| **Electron Detector** | Falcon IV |
| **Magnification** | 150,000 |
| **Voltage (kV)** | 200 |
| **Total Electron Exposure (e<sup>-</sup>/Å<sup>2</sup>)** | 60 |
| **Defocus Range (µm)** | 0.5-2.5 |
| **Pixel Size (Å)** | 0.95 |
| **Number of movie micrographs** | 2,896 |
| **Symmetry** | C1 |
| **Number of final particle images** | 77,198 |
| **Map Resolution (Å)** | 5.5 |
| **FSC Threshold** | 0.5 |
| **Sharpening Factor** | 228.3 |

